# Coping with stress: Transcriptional regulators linking cell wall integrity maintenance with primary-wall metabolism in *Arabidopsis thaliana*

**DOI:** 10.64898/2025.12.12.693917

**Authors:** Tereza Tichá, Steven Zwartkruis, Timo Engelsdorf, Zdenka Bartosova, Laura Bacete, Thorsten Hamann

## Abstract

Cell wall damage (CWD) and hyperosmotic stress elicit both distinct and overlapping transcriptional programs in plants. Responses to these stresses involve cell wall integrity (CWI) maintenance, which is mediated by receptor-like kinases such as THESEUS1 (THE1), yet the downstream regulators linking early signalling to primary cell-wall metabolism and stress-induced phytohormone biosynthesis remain poorly defined. We induced CWD and hyperosmotic stress in wild type and *the1* mutant alleles with isoxaben (ISX) and sorbitol, and analysed the gene expression of treated seedlings by RNA-seq. From these data, we focused on 15 transcriptional regulators whose expression was responsive to treatments and showed dependency on THE1 activity. Following up, we performed various functional analyses in mutant lines of these candidates and identified transcription factors that influence primary cell-wall metabolism, growth, transcriptional control of cellulose production, resistance to CWD and hyperosmotic stress, as well as phytohormone and lignin biosynthesis. Our results identify *JMJ17*, *bHLH*, and *CBP60A* as essential transcriptional regulators involved in responses to CWD and hyperosmotic stress, and they provide a starting point for dissecting the transcriptional network that regulates CWI maintenance and primary cell wall metabolism. This study sheds light on a previously unknown signalling network that regulates primary cell wall synthesis in control and stress conditions, providing a basis for more resilient crop plant breeding in the future.

**Significance Statement:** Plants provide food and materials used by society, and their productivity depends on maintaining cell wall integrity (CWI) during growth, development, and environmental stress. The CWI maintenance mechanism, involving receptor-like kinases such as THESEUS1 (THE1), has been extensively investigated, yet the transcriptional regulators involved in primary wall metabolism and stress-induced phytohormone production remain unknown. Here, we integrate transcriptomics with functional genetic and physiological assays to identify transcription factors acting in a THE1-dependent/independent manner. These regulators link CWI, hyperosmotic stress, and cell wall damage to primary wall metabolism, lignification, phytohormone production, and root growth. Our findings reveal elements of the transcriptional network that coordinate CWI and primary cell wall formation with adaptive responses, generating insights to enhance crop stress resilience.

## Introduction

The plant cell wall is a dynamic extracellular matrix that maintains cell shape and provides mechanical support during growth and development, while also contributing to resistance against biotic and abiotic stress (1). The cell wall composition and structure vary between species and tissues and are frequently remodelled during development and in response to environmental changes (2–4). The cell wall integrity (CWI) maintenance mechanism is one of the key processes activated by both abiotic and biotic stresses, as well as during development, and initiates adaptive changes in cell walls and cellular metabolism (5).

Almost all plant mature cell walls consist of primary cell walls (formed after cell division to support cell expansion) and secondary cell walls (formed after expansion has finished) (6). Secondary walls are more rigid than primary ones, providing structural support and enabling specialized functions such as water transport (7). Differences in biological function between primary and secondary cell walls are reflected in the compounds found in their respective walls (8). Primary cell walls typically contain ∼15–40% cellulose, 30–50% pectin, and 20–35% hemicellulose (such as xyloglucans or arabinoxylans), as well as cell wall proteins (8). Cellulose microfibrils confer strength and rigidity and are embedded in a pectin matrix, which is crucial for flexibility and hydration during growth. Pectins comprise homogalacturonan, rhamnogalacturonan-I (RG-I), and RG-II, which may be present in homo- or heteropolymers (3). Cell wall extensibility can be further enhanced by wall-loosening proteins such as expansins (9). Secondary cell walls are enriched in cellulose, lignin, and hemicelluloses such as xylans and glucomannans, while pectins and xyloglucans are present at much lower levels than in primary walls (10, 11). They are also less hydrated and exhibit a cellulose microfibril organisation that confers strength and resistance needed for specialised functions (e.g., woody tissue or vascular function). While lignin makes secondary cell walls more hydrophobic and has been associated with water transport (12, 13), it can also be deposited in primary cell walls in response to biotic and abiotic stress to facilitate increased stiffness and rigidity, thereby strengthening the cell wall barrier against harmful environments (such as drought or salt stress) or pathogens and pests (14–17).

The current estimate is that approximately 2,500–4,000 (∼10–15%) of the ∼27k protein-coding genes in *Arabidopsis thaliana* (called *Arabidopsis* hereafter) are involved in cell wall formation and maintenance (18, 19). Previous research has shown that transcription factors (TFs) are central coordinators regulating cell wall metabolism during growth, development, and in response to biotic/abiotic stress (20–26). Transcriptional regulation of secondary cell wall formation is well understood. Biosynthesis of the secondary cell wall is controlled through a hierarchical regulatory network involving multiple TFs that interact with each other through positive and/or negative feedback loops. This regulatory hierarchy is organized into three main tiers, each with distinct roles (11, 27). The hierarchical organisation ensures precise control over the complex process of secondary cell wall formation, which is crucial for maintaining the structural integrity of plants (3). In *Arabidopsis*, members of the NAM, ATAF, and CUC (NAC) TF family, such as SECONDARY WALL-ASSOCIATED NAC DOMAIN PROTEIN (SND), NAC SECONDARY WALL THICKENING PROMOTING FACTOR1 (NST1), and VASCULAR RELATED NAC-DOMAIN PROTEIN1 (VND1), are so-called “master regulators” because they are responsible for initiation of secondary cell wall formations in fibers (SND/NST) and vessels (VND) (28–30). The MYB-domain TFs MYB46 and MYB83 form a hub, which is essential for the regulation of cellulose, xylan, and lignin metabolism, as well as for activation of more specialized TFs (31). These more specialized TFs belong to the MYB, NAC, basic HELIX LOOP HELIX (bHLH), basic leucine zippers (bZIPs), and ETHYLENE RESPONSE FACTORS (ERF) families and are responsible for fine-tuning polymer biosynthesis and wall patterning (32). Specifically, MYB58 and -63 act as positive regulators of lignin biosynthesis while KNOTTED1-like Homeobox 7 (KNAT7) and WRKY12 modulate tissue-specific cell wall deposition (31). Influencing the expression of individual TFs has been shown to increase plant resistance to abiotic and biotic stresses (33–37).

In contrast, our understanding of the transcriptional regulation of primary cell wall synthesis and how plants modify their primary cell wall components in response to stress remains poorly understood. Currently, we have only limited information regarding the regulation of *CELLULOSE SYNTHASE* (*CesA*), which is active during primary cell wall formation (38). Cellulose synthesis in the primary cell wall is mainly conferred by CesA1, 3, and 6 (39–43). APETALA2/ETHYLENE RESPONSIVE FACTOR (AP2)/ERF Group IIId factors (ERF34/35/38/39) have been identified as specific, positive regulators of *CesA* gene expression required for primary cell wall formation (44). Interestingly, BRASSINOSTEROID INSENSITIVE1-EMS-SUPPRESSOR 1 (BES1) and BRASSINAZOLE-RESISTANT 1 (BZR1) regulate multiple CesAs, involved not only in primary cell wall synthesis but also in secondary cell wall biosynthesis (45, 46). BRASSINOSTEROID-INSENSITIVE 2 (BIN2) directly phosphorylates CesA1, linking phytohormone signalling to cellulose synthesis (45–47). The simultaneous involvement of TFs in both cell wall metabolism and stress adaptation highlights the important role of cell walls in responding to environmental stress and CWI. This also suggests that characterizing TFs involved in cell wall metabolism needs to incorporate studies investigating their roles in stress adaptation. The complex nature of the TFs network controlling secondary cell wall metabolism (11, 27) suggests that a similarly complex network could exist for primary cell wall metabolism.

Our knowledge of the transcriptional regulation of CWI maintenance mechanism is similarly limited. Plants monitor their surroundings, communicate changes in CWI, and respond with adaptive measures (5). CWI maintenance is a complex signalling network that we have only begun to understand (48–52). Cell wall damage (CWD) can occur in various ways, including during cell elongation, in response to abiotic and biotic stress, or by inhibiting cellulose production with chemicals such as isoxaben (ISX) (5, 48, 51). ISX inhibits CesA activity during primary cell wall formation, which impairs CWI and mechanical support of the cell wall, leading to severe phenotypes such as cell swelling and eventually bursting of epidermal cell walls in seedling root cells (5, 48, 51, 53). CWD induced the production of signalling molecules (among them reactive oxygen species (ROS), jasmonic acid (JA), and salicylic acid (SA)), reorganization of the cytoskeleton, changes in seedling cell wall composition (e.g. lignin deposition in primary cell walls), and mechanical properties (14, 53–55). Intriguingly, it is possible to suppress these phenotypic effects by combining ISX treatments with mild hyperosmotic stress (through simultaneous addition of osmotica such as sorbitol) (55, 56). This could be explained by the hyperosmotic stress reducing turgor pressure, which in turn alleviates the pressure on the weakened cell walls, reducing overall CWD (48). The plasma membrane-located kinases FERONIA and THESEUS1 (THE1) are required for hyperosmotic stress perception, modulation of abscisic acid (ABA) signalling, as well as CWD-induced JA and lignin production (53, 57–59). Both THE1 and FERERONIA belong to the *Catharanthus roseus* RECEPTOR-LIKE KINASE 1-LIKE kinase family (*Cr*RLK1L), whose members have been primarily implicated in biological processes involving cell wall signalling (60).

In this study, we focus on THE1 because it plays a central role in CWI maintenance and has an interesting dual function: THE1 acts as a positive regulator of CWD-induced JA and a negative regulator of hyperosmotic stress-induced ABA production, thereby modulating the mechanical properties of plant cell walls (53). There is limited information about interaction partners of THE1, although it is known that THE1 directly interacts with GUANINE EXCHANGE FACTOR 4 (GEF4) (61). GEFs can activate GTP-hydrolyzing proteins of the Rho-related GTPases of Plants (ROP) family, which act as molecular switches and signal transduction proteins (62). This interaction regulates other components through phosphorylation, and some of their targets include RESPIRATORY BURST OXIDASE HOMOLOG D (RBOHD) and calcium channels, linking receptor activation and secondary responses (63–65).

This study investigates transcriptional regulators downstream of THE1 and connections between CWI maintenance and primary cell wall metabolism. We aim to address knowledge gaps of the transcriptional regulation of CWI maintenance and primary cell wall metabolism through an integrated experimental approach. We perform RNAseq-based expression studies of wildtype and *the1* mutant seedlings exposed to ISX and hyperosmotic stress treatments. We use the resulting expression data to identify 15 candidate genes encoding putative transcriptional regulators (called hereafter candidates) that exhibit THE1-dependent or -independent expression changes upon exposure to ISX-induced CWD or sorbitol-induced hyperosmotic stress. We investigate the biological functions of the 15 candidates by selecting T-DNA insertion alleles, determining the impact of the insertions on cell wall composition and root growth in a hypersensitivity screen, assessing the impact of the candidates on stress-induced phytohormone and lignin production, and establishing the roles of the candidates in primary cell wall synthesis and transcriptional regulation of *CesA* genes. Our results identify putative transcriptional regulators required for CWI maintenance as well as primary cell wall metabolism and show that the expression of specific candidate genes is regulated by THE1 activity.

## Results

### Identification of candidate transcription factors mediating THE1-based cell wall integrity signaling

THE1 acts as a positive regulator of CWD-induced JA and a negative regulator of hyperosmotic stress-induced ABA production while modulating the mechanical properties of cell walls (66). Here we aimed to identify transcriptional regulators of primary cell wall biosynthesis that act in a THE1-dependent manner and in response to hyper-osmotic stress or CWD by combining transcriptomic with functional studies (Fig. 1A). To achieve this, we exposed 7-day-old Arabidopsis seedlings for 2 hours to sorbitol (inducing hyperosmotic stress), isoxaben (ISX), combined sorbitol/ISX, or control (DMSO) conditions. At this developmental age, the seedlings have normally formed very little secondary cell walls, thus ensuring we are primarily studying processes related to primary cell wall metabolism (7). The stress conditions were selected based on previously established experimental protocols (53). We employed the following genotypes: Col-0, *the1-1* (loss-of-function allele (67)), and *the1-4* (hypermorphic allele (68)). RNA-seq analysis of the samples revealed that sorbitol and sorbitol/ISX treatments induced the most profound changes in transcript levels, based on the number of differentially expressed genes (DEGs; Fig. 1B). Combining individual sorbitol and ISX treatments had a very limited additive effect on the number of DEGs (approximately 2% more than sorbitol alone), with approximately 80% of the same genes being affected by both treatments. Transcriptional changes were similar between Col-0 and *the1-1*, whereas *the1-4* seedlings exhibited 25% more DEGs after ISX treatment than Col-0. Gene Ontology (GO) enrichment analysis of the RNA-seq data with respect to the molecular function revealed that the strongest enrichment of DEGs was associated with transcriptional regulation and DNA-binding transcription factor activity (Fig. 1C). GO analysis of biological processes showed strong enrichment for responses to stress, chemical, and abiotic stimuli (SI Appendix, Fig. S1A). In the cellular component GO category, DEGs primarily belonged to the cell periphery and plasma membrane (SI Appendix, Fig. S1B). To identify relevant candidate genes, we applied two selection criteria to the RNA-seq dataset. First, genes had to have predicted DNA-binding transcription factor activity (GO:0003700). This criterion was met by 8% of all DEGs (478 out of 5,826). Second, we prioritized genes whose expression changed in response to ISX treatments and was influenced by *THE1* activity. Within this group, those showing opposite expression patterns in *the1-1* and *the1-4* seedlings were considered especially interesting (Fig. 1D, SI Appendix, Fig. S2). Based on these criteria, we selected 15 DEGs for follow-up studies. Several of these DEGs have been shown before to act as transcription factors (TFs) in processes such as plant adaptation to light, cell cycle progression, regulation of secondary cell wall metabolism, or ABA production in response to nitrate levels and seed maturation (69–76). The underlying common denominator of these processes being active cell wall remodeling taking place.

**Figure 1.**
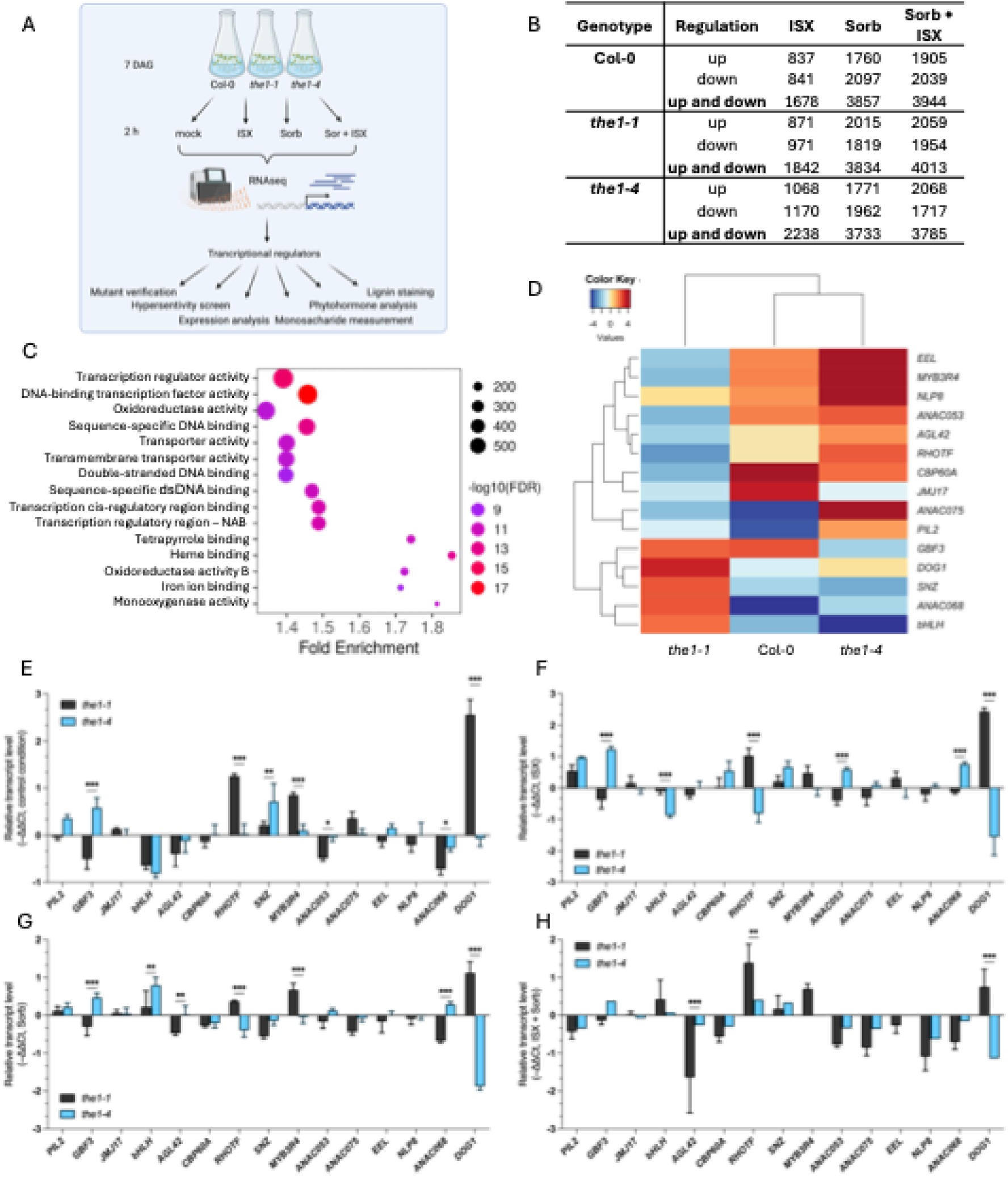
Transcriptomic RNA-seq analysis of Arabidopsis genotypes in response to ISX and osmotic stress. (A) Schematic overview of the experimental setup; Created in https://BioRender.com. (B) Table summarizing up- and downregulated differentially expressed (DE) genes in 7-day-old Arabidopsis seedlings following exposure to sorbitol, ISX, and combined stress, relative to the mock condition. Genes were considered differentially expressed if they met the criteria: FDR < 0.05 and |log_2_FC| > 0.585. (C) Gene Ontology (GO) enrichment analysis for Molecular Function (GO-MF) based on DE genes identified in Col-0, *the1-1*, and *the1-4* under ISX, sorbitol, and combined ISX + sorbitol treatments. The top 15 GO-MF terms (FDR < 0.05) are displayed. The X-axis represents fold enrichment of each GO term. The size of each circle corresponds to the number of genes associated with that term, and the color gradient from purple to red indicates –log₁₀(FDR), with red representing higher statistical significance. Analysis was performed using ShinyGO v0.82 (Ge SX, Jung D & Yao R, Bioinformatics 36:2628–2629, 2020). Abbreviations: Oxidoreductase activity B – Oxidoreductase activity acting on paired donors with incorporation or reduction of molecular oxygen; Transcription regulatory region – NAB – Transcription regulatory region nucleic acid binding; dsDNA binding – Sequence-specific double-stranded DNA binding. (D) Heatmap showing expression levels of selected candidate genes under 600 nM ISX treatment in Col-0, *the1-1*, and *the1-4*. Heatmap color scaling is based on relative fold change, with thresholds set between –4.5 and 4.5. Hierarchical clustering using Euclidean distance and complete linkage is shown as dendrograms for both rows and columns. (E–H) Expression levels of selected candidate genes in *the1-1*, and *the1-4* seedlings after 6 h of treatment (E) Control (DMSO), (F) 600 nM Isoxaben (ISX), (G) 300 mM Sorbitol, and (H) combined ISX + Sorbitol. Transcript levels were normalized to *SEC3a* and are presented as –ΔΔCt values (log_2_-transformed relative expression) compared to mock-treated controls within each genotype. Data represent mean ± SD (n ≥ 3) from three independent experiments. Asterisks indicate statistically significant differences between *the1-1* and *the1-4* and their respective controls (two-way ANOVA followed by Šídák’s multiple comparisons test; *p < 0.05; **p < 0.01; ***p < 0.001).

### Investigating the expression patterns of candidate genes

To investigate the expression dynamics of the selected candidate genes and their dependency on THE1, we performed quantitative reverse transcription PCR (qRT-PCR)-based expression analysis in Col-0, *the1-1*, and *the1-4* seedlings after 6 hours of mock, ISX, sorbitol, and sorbitol/ISX treatments (Fig. 1E-H, SI Appendix, Fig. S3). ISX treatments induced limited transcriptional changes in candidate genes in Col-0 seedlings. Gene expression of *CBP60A* was increased, while *SNZ* expression was decreased (SI Appendix, Fig. S3A). In contrast, gene expression of 10 out of 15 candidate genes was affected in response to sorbitol (no pronounced expression changes observed in *JMJ17*, *AGL42*, *CBP60A*, *EEL,* and *ANAC068*). Expression of 11 candidates was affected by sorbitol/ISX co-treatments (not *JMJ17*, *AGL42*, *RHOTF*, and *EEL*). In ISX-treated *the1-1* seedlings, expression of *AGL42*, *CBP60A*, and *EEL* was increased, while *SNZ* and *ANAC075* expression was decreased (SI Appendix, Fig. S3B). In response to sorbitol, expression increased in *PIL2*, *GBF3*, *ANAC053*, and *NLP8*, while *bHLH* and *MYB3R4* expression was decreased. In response to sorbitol/ISX co-treatments, *PIL2*, *GBF3*, *SNZ*, *ANAC053*, *NLP8*, and *ANAC068* expression was increased, whereas *bHLH*, *AGL42*, and *MYB3R4* expression was decreased. In ISX-treated *the1-4* seedlings, expression of *GBF3*, *AGL42*, *CBP60A*, *ANAC053*, *EEL*, and *ANAC068* increased, and *RHOTF*, *SNZ*, and *DOG1* expression decreased (SI Appendix, Fig. S3C). Expression of *PIL2*, *GBF3*, *RHOTF*, *SNZ*, *ANAC053*, *ANAC075*, and *NLP8* was increased in sorbitol-treated seedlings, and *bHLH* and *MYB3R4* expression was decreased. Upon sorbitol/ISX co-treatments, *PIL2*, *GBF3*, *SNZ*, *ANAC053*, *ANAC075*, *NLP8*, *ANAC068*, and *DOG1* expression was increased, while *bHLH*, *AGL42*, and *MYB3R4* expression was decreased. To facilitate understanding of the function of *THE1* in regulating candidate gene expression, we replotted the qRT-PCR-derived expression data (Fig. 1E-H). The candidate genes can be divided into three different groups based on their expression patterns. Group 1 consists of candidate genes whose expression is independent of *THE1* activity across all tested conditions. *PIL2*, *JMJ17*, *CBP60A*, *ANAC075*, *EEL*, and *NLP8* belong to this group. Group 2 consists of candidates whose gene expression is condition-specifically dependent on *THE1* activity. *GBF3*, *ANAC068*, *AGL42*, *ANAC053*, *SNZ*, and *MYB3R4* form this group. Group 3 consists of *DOG1* and *RHOTF*, as their expression is consistently dependent on *THE1* activity across all treatments. qRT-PCR-based expression analysis after 6 hours revealed similar but not identical expression patterns for the candidate genes compared to the RNA-seq data (2 hour time point), possibly reflecting differences between the early and later phases of transcriptional responses to stress treatments. 11 of 15 candidates show expression changes in response to treatments, but only *DOG1* and *RHOTF* show consistent, *THE1*-dependent, transcriptional changes.

### Assessing the biological relevance of candidate genes

We isolated homozygous T-DNA insertion lines and investigated the impact of the insertions on candidate gene expression by qRT-PCR (SI Appendix, Table 1, and SI Appendix, Fig. S4). We identified 12 lines representing knock-out (KO), two knock-down (KD; *pil2* and *agl42*), and overexpression (OE; *anac068*) alleles. Initially, we performed a hypersensitivity assay using root length as a readout to determine the biological relevance of candidate genes in the response to CWD and hyperosmotic stress. Candidate insertion line and wild-type seedlings were grown on plates with ½ MS medium (control) or plates supplemented with either 200 mM sorbitol or 2 nM ISX (Fig. 2, SI Appendix, Fig. S5). *anac068*, *agl42*, *gbf3, bhlh, cbp60a, snz, anac053, anac075, eel, nlp8,* and *dog1* seedlings exhibited increased root growth in control condition. In contrast, *rhotf* seedlings displayed reduced root growth. Col-0 seedlings exhibited more than 50% reduction in root length in response to ISX compared to control conditions. *jmj17*, *snz*, *myb3r4*, *eel*, *nlp8*, and *dog1* seedlings showed hypersensitive root growth on ISX-containing medium. Col-0 seedlings showed approximately 40% reduction in root growth in response to sorbitol treatments. While *pil2* and *nlp8* seedling roots were hyposensitive to sorbitol, *jmj17* and *eel* were hypersensitive. The results of the phenotypic analysis suggested that *GBF3, BHLH, CBP60A, SNZ, ANAC053, ANAC075, EEL, AGL42, NLP8,* and *DOG1* act as negative regulators of root growth, while *ANAC068* and *RHOTF* act as positive ones. Based on stress-specific root growth phenotypes, the candidate insertion lines can be grouped into three categories. The first category, where root growth is reduced on ISX, includes *snz*, *myb3r4*, and *dog1*, suggesting that *SNZ*, *MYB3R4,* and *DOG1* act as positive regulators of root growth in response to ISX. The second category, where root growth is affected by sorbitol, comprises only *PIL2*, which appears to act as a negative regulator of root growth in response to sorbitol treatments. The last category, where root growth is affected by both sorbitol and ISX, includes *jmj17*, *eel*, and *nlp8*. *NLP8* seems to act as a positive regulator of root growth in response to ISX and a negative regulator to sorbitol, whereas *JMJ17* and *EEL* are positive regulators in response to both ISX and sorbitol. The results of the hypersensitivity screen suggest that three of the identified candidates (*PIL2*, *JMJ17,* and *MYB3R4*) have specific functions in responses to CWD and/or hyper-osmotic stress, while the remaining candidates are required for general growth-related processes.

**Figure 2.**
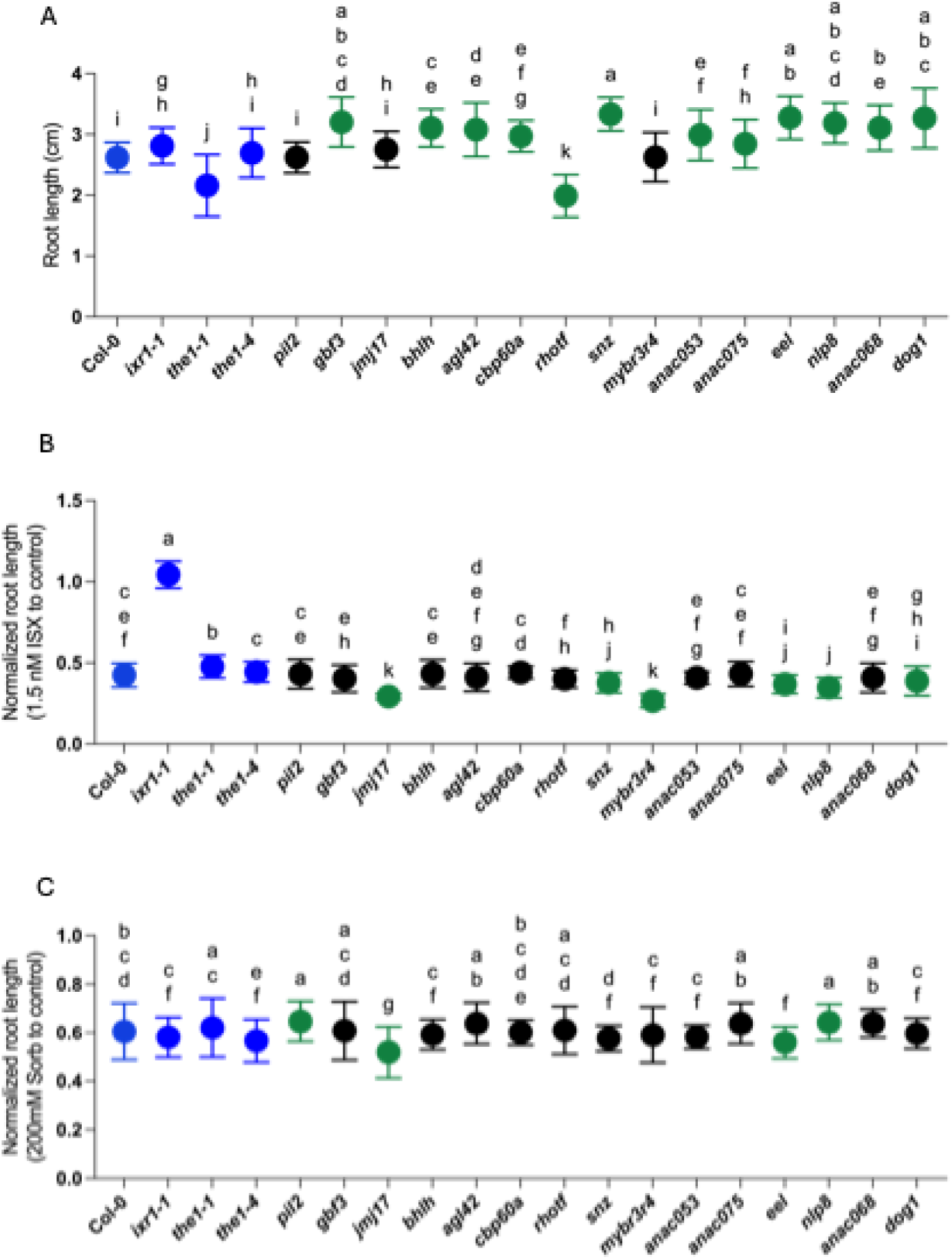
Hypersensitivity assay based on root length measurements of Arabidopsis seedlings grown under control and stress conditions. (A) Root length of 7-day-old seedlings grown on control medium (½ MS). (B) Relative root length of seedlings grown on 1.5 nM Isoxaben (ISX), normalized to control conditions. (C) Relative root length of seedlings grown on 200 mM sorbitol, normalized to control conditions. Data represent mean ± SD (n ≥ 120) from three independent experiments. Different letters indicate statistically significant differences between genotypes (one-way ANOVA followed by Tukey’s HSD test; p < 0.05). Genotypes significantly different from Col-0 are highlighted in green.

### Analysis of cell wall composition

To determine whether the candidates influence primary cell wall composition, we quantified monosaccharide composition of cell walls for 7-day-old seedlings of the respective candidate mutant lines (Fig. 3, SI Appendix, Fig. S6, S7). No changes in glucuronic acid levels were observed in any of the genotypes examined. However, alterations in other monosaccharides were detected (Fig. 3A–I). Fucose levels were increased in *jmj17*, *agl42*, *eel*, and *nlp8* seedlings, while rhamnose levels were increased in *pil2*, *snz*, *myb3r4*, *anac053*, and *dog1*. Arabinose was increased in *myb3r4* and *eel* seedlings, and galactose increased in *rhotf*, *snz*, *myb3r4*, and *dog1*. Xylose levels were reduced only in *rhotf* seedlings. Mannose increased in seedlings of *rhotf*, *snz*, and *anac075*. Galacturonic acid levels were elevated in *anac075* seedlings, and decreased in *jmj17*, *rhotf*, *myb3r4*, *eel*, *nlp8*, and *dog1*. Interestingly, cellulose was most affected in the seedlings examined, with significantly reduced levels observed in 9 out of 15 genotypes examined (*pil2*, *gbf3*, *bhlh*, *cbp60a*, *rhotf*, *snz*, *myb3r4*, *eel*, and *dog1).* In contrast, no significant differences were observed in *the1-1* and *the1-4* seedlings (SI Appendix, Fig. S6). We performed a phenotypic clustering using the results from the cell wall analysis to determine if candidates are modulating particular aspects of cell wall metabolism jointly (Fig. 3J). *bhlh*, *dog1*, and *rhotf* seedlings showed similar effects on glucuronic acid, galacturonic acid, and glucose content, suggesting a shared role in pathways related to cellulose, hemicellulose, and pectin biosynthesis. Furthermore, *dog1* and *rhotf* seedlings displayed similar effects on galactose, rhamnose, and mannose levels, sugars mainly associated with pectin (galactose and rhamnose), glycoproteins (galactose and mannose), and hemicellulose (mannose). *jmj17*, *eel*, and *nlp8* seedlings exhibited similar effects on xylose and fucose levels, key components of hemicellulose and pectin. Several mutants, including *dog1*, *rhotf*, *eel*, *snz*, *myb3r4*, *pil2*, *nlp8*, *anac075*, and *jmj17,* exhibited significant differences in two or more types of monosaccharides. We summarized the results from the monosaccharide analysis (SI Appendix, Fig. S7). Interestingly, many of the selected candidate genes (*JMJ17, EEL, PIL2, SNZ, MYB3R4, RHOTF, GBF3, bHLH, CBP60A,* and *DOG1*) seem to act as positive regulators of cellulose biosynthesis. Mutant seedlings for *RHOTF* and *DOG1*, which are apparently regulated by THE1, exhibit similar cell wall defects. Our results implicate 14 of the 15 candidates characterized in the regulation of primary cell wall metabolism.

**Figure 3.**
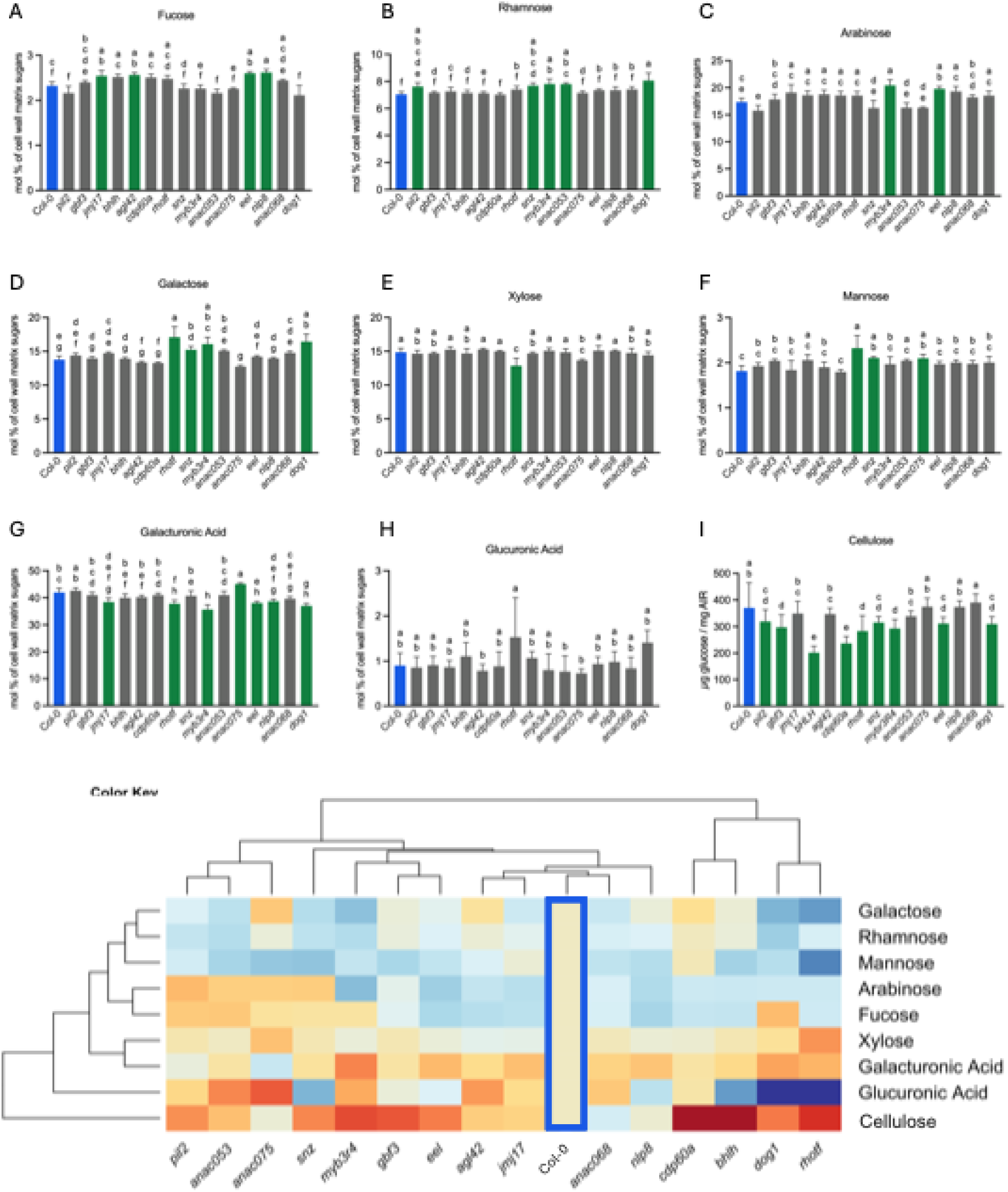
Analysis of cell wall components in 7-day-old Arabidopsis seedlings. (A–H) Molar percentage (mol%) of individual matrix polysaccharide sugars ((A) fucose, (B) rhamnose, (C) arabinose, (D) galactose, (E) xylose, (F) mannose, (G) galacturonic acid, and (H) glucuronic acid) in the indicated genotypes, derived from alcohol-insoluble residue (AIR). (I) Cellulose content (µg glucose / mg AIR) measured using the anthrone assay. Data represent mean ± SD from five independent biological replicates. Statistical significance was determined by one-way ANOVA followed by Tukey’s HSD test (p < 0.05). Different letters indicate statistically significant differences between genotypes. Green-colored bars indicate significant differences compared to Col-0. (J) Heatmap showing log_2_-transformed fold-change values for each sugar in mutant lines relative to Col-0 (mutant/Col-0). Hierarchical clustering using Euclidean distance and complete linkage is shown as dendrograms for both rows and columns. The color scale is restricted to -0.5 to +0.5 to emphasize moderate differences in sugar levels.

### Transcriptional regulation of primary cell wall *CesA* genes by ISX and osmotic stress

Next, we determined if expression of the primary cell wall cellulose synthase genes (*CesAs*; *CesA1*, *CesA3*, and *CesA6*) is changing in response to CWD and osmotic stress in Col-0, *the1-1*, *the1-4*, and candidate mutant seedlings after a 6-hour treatment with either ISX or sorbitol using qRT-PCR (Fig. 4, and SI Appendix, Fig. S8, S9). RNA-seq-based expression analysis had shown that expression of *CesA1*, *3*, and *6* is decreased in sorbitol and sorbitol/ISX-treated Col-0 seedlings after two hours, while ISX treatments seemed to have qualitatively different effects (SI Appendix, Fig. S8). qRT-PCR-based expression analysis of *CesA1*, *3*, and *6* in mock-, ISX- or sorbitol-treated Col-0 seedlings detected similarly decreased expression in response to sorbitol treatments for all three genes (Fig. 4A). Furthermore, expression of *CesA1*, *3,* and *6* in *the1-1* and *the1-4* seedlings showed no significant differences in control conditions to Col-0 (SI Appendix, Fig. S9). In ISX-treated seedlings, *CesA1* expression was decreased in *the1-4* compared to the relevant control. In sorbitol-treated seedlings, *CesA1* and *CesA6* expression were decreased in *the1-1* compared to Col-0. Next, we determined *CesA1*, *CesA3*, and *CesA6* expression in ISX or sorbitol-treated candidate mutant seedlings (Fig. 4B, C). 12 of 15 ISX-treated candidate lines showed no significant differences in expression of *CesA1, 3,* and *6* compared to Col-0. However, *bhlh* and *cbp60a* seedlings exhibited significantly increased *CesA1* expression, while *nlp8* seedlings showed decreased *CesA3* and *CesA6* expression. Sorbitol-treated candidate seedlings exhibited reduced *CesA1* expression in 13 of 15 tested alleles compared to sorbitol-treated Col-0 (exceptions being *snz* and *anac068*; Fig. 4C). Interestingly, *CesA3* expression was increased in *snz* and *anac068* and decreased in *anac075* seedlings compared to Col-0. *CesA6* expression was increased in *snz* seedlings and decreased in *gbf3*, *anac053*, and *anac075* seedlings relative to Col-0. To summarize, qRT-PCR-based expression analysis revealed that osmotic stress and ISX-induced CWD have qualitatively different effects on expression of *CesA1*, *CesA3*, and *CesA6* in Col-0 seedlings. Gene expression data from ISX-treated candidate seedlings suggests that *bHLH* and *CBP60A* function as negative regulators of *CesA1* expression, whereas *NLP8* apparently regulates positively the expression of *CesA3* and *CesA6*. Results from sorbitol-treated candidate insertion lines suggest that *PIL2*, *GBF3*, *JMJ17*, *bHLH*, *AGL42*, *CBP60A*, *RHOTF*, *ANAC053*, *ANAC075*, *EEL, NLP8*, and *DOG1* may act as positive regulators of *CesA1* expression. *SNZ* seems to function as a negative regulator of *CesA3* expression, whereas *ANAC075* and *ANAC068* seem to act as positive regulators. *GBF3*, *ANAC053*, and *ANAC075* seemingly act as positive regulators of *CesA6*. Interestingly, the results for three candidate genes suggest they may have a stress-specific dual role in response to CWD and hyperosmotic stress. *bHLH* and *CBP60A* seem to act as positive regulators of *CesA1* expression in response to sorbitol but as negative regulators in response to ISX. *NLP8* acts as a positive regulator for both *CesA3* and *CesA6* expression in response to ISX, and *CesA1* expression in response to sorbitol. These findings suggest that the majority of our candidates are involved in the transcriptional regulation of *CesA* genes, which are involved in primary cell wall formation, upon exposure to CWD and/or hyperosmotic stress.

**Figure 4.**
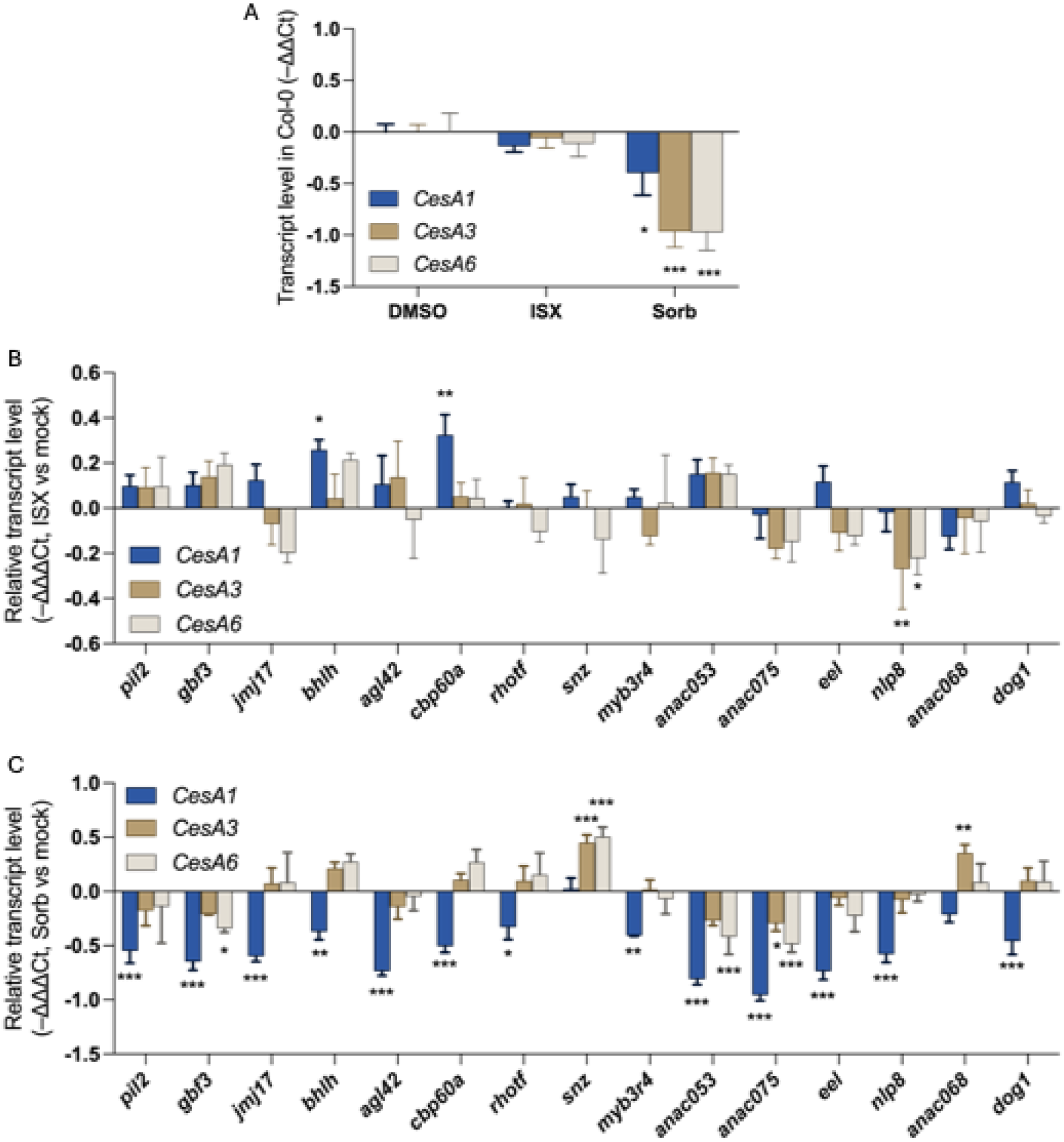
Quantitative RT-PCR analysis of *CesA* genes and selected candidate genes in Arabidopsis seedlings in control and stress conditions. (A–C) Transcript levels of *CesA1*, *CesA3*, and *CesA6* in Col-0 and candidate seedlings after 6 h treatment with control (fresh media), 600 nM Isoxaben (ISX), or 300 mM sorbitol (Sorb). (A) *CesA1*, *CesA3*, and *CesA6* transcript levels in Col-0 under DMSO, ISX, and Sorb treatments. (B, C) Relative transcript levels in indicated genotypes following (B) 600 nM ISX or (C) 300 mM Sorbitol treatment. Transcript levels were normalized to SEC3a and presented as –ΔΔCt values (log_2_-transformed relative expression), with additional normalization to mock-treated Col-0. For panels and (C), values were first normalized to mock-treated samples within each genotype, then to Col-0, and are shown as –ΔΔΔCt values. Bars represent mean ± SD (n ≥ 3) from three independent biological replicates. Asterisks indicate statistically significant differences compared to Col-0, calculated from ΔΔCt values (two-way ANOVA followed by Dunnett’s post hoc test; *p < 0.05; **p < 0.01; ***p < 0.001).

### Phytohormone and lignin production in candidate mutant seedlings

To determine whether the candidates are involved in established responses to CWD and hyperosmotic stress, we performed quantitative analysis of JA, ABA, and lignin levels in candidate mutant seedlings treated with ISX, sorbitol, or sorbitol/ISX (Fig. 5, SI Appendix, Fig. S10-S13) (55, 66). In control conditions, ABA and JA levels were at the detection limit (SI Appendix, Fig. S10A, S10B). After 4 hours of sorbitol treatment, ABA production was similarly induced in Col-0 and all candidate seedlings (Fig. 5A). ISX-induced production of JA after 6 hours was observed in all genotypes, with *pil2*, *jmj17*, *rhotf*, and *anac075* seedlings showing significantly higher JA levels than Col-0 (Fig. 5B). After 4 hours of combined sorbitol/ISX treatment, ABA accumulation was similar to the responses seen with sorbitol alone (SI Appendix, Fig. S10C). The combined treatments did not induce JA production in any genotype compared to the control (SI Appendix, Fig. S10C and S10D). Next, we quantified CWD-induced lignin deposition in seedling root tip areas using an image analysis-based approach (55). Lignin deposition was visualized using phloroglucinol staining in Col-0 and candidate seedlings (Fig. 5C, SI Appendix, Fig. S11, S12). Image-based analysis of the lignified area revealed that *the1-1* seedlings had significantly reduced lignin production, consistent with the previously established function of THE1 in CWI maintenance (SI Appendix, Fig. S13; (67)). *pil2*, *jmj17*, *rhotf*, and *anac075* seedlings also exhibited significantly reduced lignification, suggesting impaired activation of lignin production. To summarize, these results suggest that none of the candidates is required for sorbitol-induced ABA production. In contrast, *PIL2*, *JMJ17*, *RHOTF,* and *ANAC075* serve as negative regulators of CWD-induced JA production, and *PIL2*, *JMJ17*, *GBF3*, *SNZ*, and *NLP8* seem to act as positive regulators of lignin deposition in response to CWD. These findings suggest that several of the candidates investigated here are involved in JA and lignin production, implicating them in CWI maintenance.

**Figure 5.**
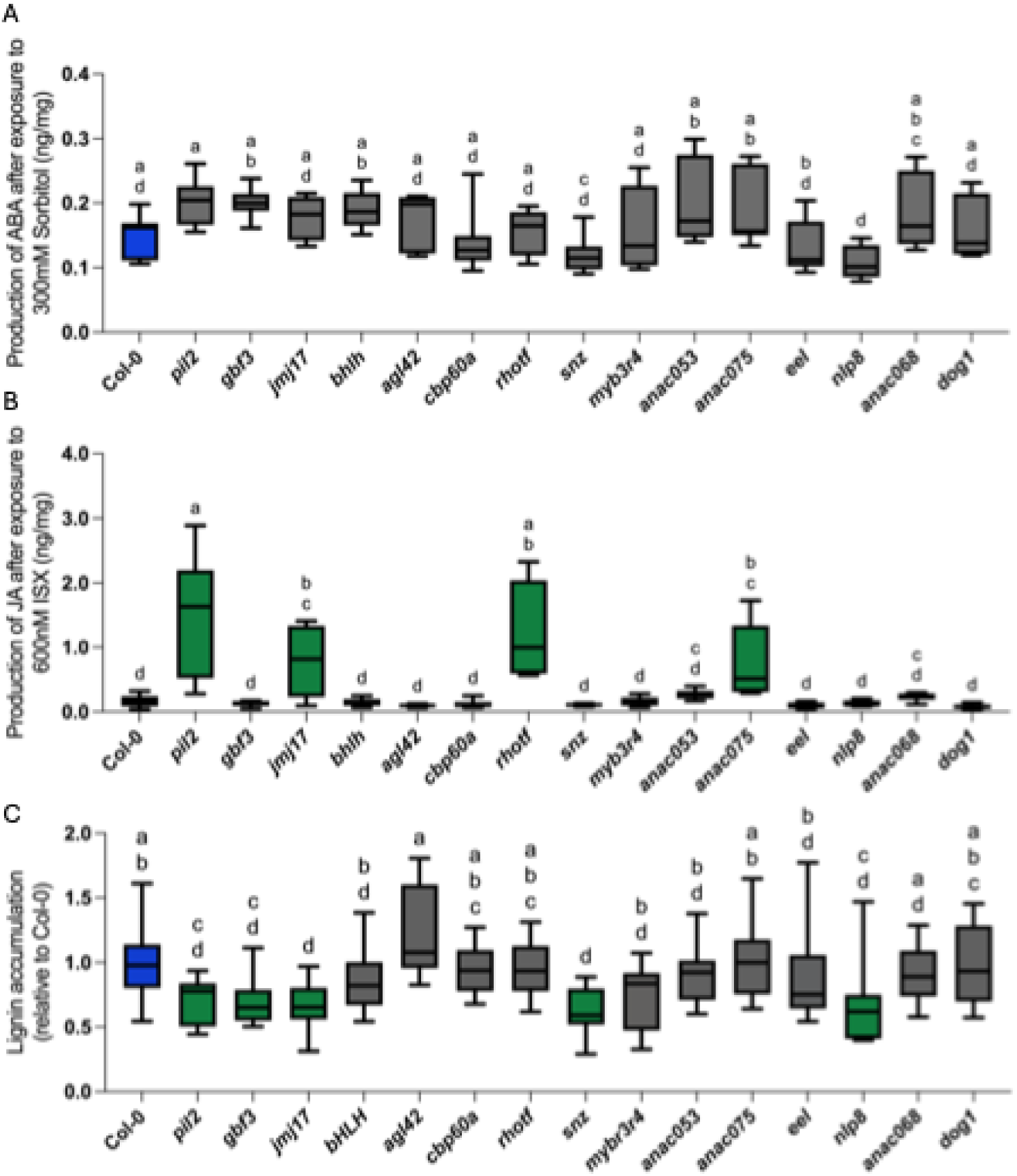
Jasmonic acid, abscisic acid, and lignin accumulation in candidate T-DNA insertion mutants under stress conditions. (A) Abscisic acid (nanograms/milligrams dry weight) levels after 4 h of 300 mM sorbitol treatment. (B) Jasmonic acid (nanogram/milligram dry weight) levels after 6 h of 600 nM Isoxaben (ISX) treatment. (C) Relative lignin integrated density in root tips after 12 h of ISX treatment, expressed as mutant/Col-0 ratio from the measured stained area by the phloroglucinol assay. Graphs represent data from 3 to 4 independent experiments. Box plots display the full range (from minimum to maximum), with horizontal bars indicating the mean. Different letters indicate statistically significant differences between genotypes (one-way ANOVA followed by Tukey’s HSD test; *p < 0.05). Green-colored box plots indicate significant differences compared to Col-0-treated seedlings.

## Discussion

Here, we aimed to characterize the THE1-dependent transcriptional network modulating stress responses in seedlings exposed to CWD and sorbitol-induced hyperosmotic stress, and to explore whether the same network also regulates primary cell wall metabolism. We performed RNA-seq analysis of Col-0, *the1-1,* and *the1-4* seedlings exposed to CWD (ISX) or hyperosmotic stress (sorbitol) and selected 15 candidates for follow-up studies. Based on previously published results, 12 of the selected candidates are TFs, while *DOG1*, *RHOTF*, and *JMJ17* are not typical TFs but have been implicated in the transcriptional regulation of plant stress responses (77–80). We isolated KO/KD/OX alleles for the candidates to perform targeted phenotypic analyses, including growth and stress assays, as well as cell wall, phytohormones, and gene expression analyses.

Our RNA-seq–based gene expression analysis showed that sorbitol and sorbitol/ISX treatments elicited significantly more DEGs than ISX alone, with the two sorbitol-containing treatments having very similar effects. This is in line with our current understanding that hyper-osmotic stress triggers global, ABA-mediated adaptive responses, while ISX-treatments have, in the short term, limited effects (1, 81–86). A notable fraction of the DEGs encodes transcriptional regulators, probably responsible for transcriptional reprogramming induced by ISX and/or sorbitol treatment initiated within hours after starting the treatment. THE1 mediated responses to ISX involve ROS, JA, and SA production, nitrate metabolism, cell cycle arrest involving cytokinin-based signaling, as well as cell wall modifications such as lignification, while hyper-osmotic stress increases ABA levels and induces adaptive changes in growth (14, 55, 66, 87–91). The 15 candidates were selected based on a combination of criteria, including significant changes in transcript levels in ISX-, sorbitol-, or sorbitol/ISX-treated seedlings. qRT-PCR-based expression analysis of the same genes in the same genotypes (Col-0, *the1-1*, *the1-4*) and treatments after 6 hours (Fig. 1E-H, SI Appendix, Fig. S3) revealed similarities and differences compared to the RNAseq-derived data obtained after 2 hours of treatment (Fig. 1D, SI Appendix, Fig. S2). We consider the differences indicative of dynamic transcriptional changes during the early phases of exposure to CWD and hyperosmotic stress. Based on the expression analysis results (Fig. 1E-H), we assigned the candidates to three groups. Group I (*PIL2/PIF6*, *JMJ17*, *CBP60A*, *ANAC075, EEL/ATBZIP12, bHLH,* and *NLP8*) consisted of candidates exhibiting THE1-independent, stress-induced expression changes. Previous work has implicated the candidates in adaptation to light (*PIL2/PIF6* (*92*), *JMJ17* (*78, 79*)), regulation of ROS levels during senescence (*ANAC075* (*71*)), ABA-mediated processes involving ABI5 (*NLP8* (*93*)), *EEL/ATBZIP12* (*94*)), or salicylic acid related to plant immunity (*CBDP60A* (*95*)). Group II (*GBF3*, *ANAC068*, *AGL42*, *ANAC053*, *SNZ*, and *MYB3R4*) candidates exhibited condition-specific, THE1-dependent changes. These TFs have been implicated in modulation of ROS levels (*AGL42* (*96*)), cell cycle regulation (*MYB3R4* (*76*), *ANAC068* (*97*)), response to ABA (*GBF3* (*98, 99*)), and modulation of apoplastic immunity, possibly via regulation of xyloglucan metabolism (*ANAC053* (*100, 101*)) or floral transition (*SNZ* (*102*)). Group III consists of *DOG1* (*80*) and *RHOTF,* which exhibited THE1-dependent expression changes across all conditions and timepoints examined. Currently, information on the biological processes, for which *RHOTF* is required, is limited (103). Protein homology searches using the RHOTF sequence identify a domain at the C-terminus with similarity to RHO-N-1, a transcription terminator exhibiting RNA-dependent ATPase activity (104). *DOG1* has been previously implicated in regulating seed germination in response to temperature and ABA-based signaling, a process that conceivably can involve cell wall metabolism and impair CWI (80). Similar considerations apply to all the other candidates identified. They may mediate cell wall-related aspects of the biological processes they have been previously implicated in.

We tested this hypothesis by performing cell wall analysis of mock-treated seven-day-old mutant seedlings, which consist mostly of primary cell walls. The results of the analysis indicate that 14 of the 15 genotypes examined exhibit significant changes often in more than one cell wall component (exception *anac068*, Fig. 3, and SI Appendix, Fig. S7). Intriguingly, *anac075*, *jmj17*, *eel*, *myb3r4*, *rhotf*, *nlp8* and *dog1* seedlings exhibited significant differences in galacturonic acid levels, implicating them in the regulation of pectin metabolism. *jmj17*, *eel*, *myb3r4*, *snz*, *rhotf*, *dog1*, *pil2*, *gbf3*, *bhlh,* and *cbp60a* seedlings exhibited reduced cellulose levels, suggesting that these TFs act as positive regulators of cellulose biosynthesis. Considering these results and the fact that the sample material consists mostly of primary cell walls, it is reasonable to conclude that the candidates identified here are regulating different aspects of primary cell wall metabolism.

Primary-wall cellulose synthesis in Arabidopsis relies on CesA1, CesA3, and CesA6 (39–43). Previous work showed that abiotic stress leads to reduced cellulose levels either by changes in *CesA* expression and/or by altering cellulose synthase complex localization (1, 85, 86, 105–107). ISX-induced CWD normally acts on the post-transcriptional level by changing the localization or activity of cellulose synthase complexes (47, 55, 67, 89, 105, 108–110). Consistently, we observed reduced expression of *CesA1/3/6* in sorbitol-treated Col-0 seedlings after six hours, whereas ISX treatments had no significant effects on transcript levels (Fig. 4A). When investigating *CesA1/3/6* expression in mutant seedlings, sorbitol- or ISX-treated, we observed altered gene expression (Fig. 4B, C). The results suggest that *bHLH*, *CBP60A,* and *NLP8* modulate *CesA1/3/6* expression in response to ISX treatment, but it remains to be determined how direct or indirect they act. All sorbitol-treated mutant seedlings exhibited differences from the corresponding Col-0 controls for at least one of the *CesAs* investigated. For 12 alleles, the effects were qualitatively similar to the sorbitol effects on *CesA1/3/6* expression observed in Col-0, suggesting that they modulate *CesA* expression in response to hyper-osmotic stress. However, in *snz* and *anac068* seedlings, the effects were more pronounced, suggesting that the TFs play a fundamental role in regulating *CesA1/3/6* expression. Previously, we postulated the existence of a mechanism that modulates adaptive responses (such as changes in cell wall metabolism) in plants by integrating signals caused by CWD and hyperosmotic stress (48). Such a mechanism would require TFs regulating cell wall metabolism and the transcriptional response to both ISX and sorbitol treatments. Based on the results of RNAseq/qRT-PCR expression and cell wall analyses, *bHLH* and *CBP60A* seem to meet these criteria (Fig. 4B, C, and SI Appendix, Fig. S7).

Seedlings for 12 of the 15 mutant alleles examined exhibit seedling root growth defects (Fig. 2A), suggesting the genes affected are required for root growth. Several of the mutant seedling roots also exhibit hypersensitivity/resistance phenotypes when continuously exposed to ISX or sorbitol (Fig. 2B, C), suggesting that they are required for growth-related processes, particularly affected by these stresses. Interestingly, *jmj17*, *pil2,* and *myb3r4* seedlings do not exhibit growth defects in control condition but are either more resistant to ISX (*pil2*), more sensitive to hyperosmotic stress (*myb3r4*), or more sensitive to both stresses (*jmj17*). These results implicate *PIL2* and *MYB3R4* in the specific response to a particular stress, whereas *JMJ17* appears to mediate both responses. The results from the phytohormone and lignification assays indicate that *MYB3R4* is not required for ISX- or sorbitol-induced responses. However, both *PIL2* and *JMJ17* are negative regulators of ISX-induced JA production and positive regulators of lignification (Fig. 5B, C). These results are in accordance with previous data showing that seedlings deficient in ALLENE OXIDE SYNTHASE (AOS, an enzyme that gives rise to the JA precursor 12-oxophytodienoic acid) have elevated lignification (14). However, elevated JA levels alone seem insufficient to repress lignification, since *rhotf* and *anac075* seedlings exhibit elevated JA levels but normal lignification. Furthermore, *gbf3, snz,* and *nlp8* seedlings exhibit reduced lignification while JA levels are normal, further supporting the notion that JA induction and lignification are two separate CWD-induced responses. The results from the stress assays suggest that *GBF3, SNZ,* and *NLP8* are required for CWD-induced lignification and processes underlying root growth in general. It is intriguing to note that *eel* and *nlp8* exhibited increased resistance to sorbitol, while *jmj17* and *pil2* seedlings showed increased sensitivity, and ABA levels remained normal compared to wild type. This suggests that the modified response to hyperosmotic stress involves an ABA-independent process. The results from the growth, phytohormone, and lignification assays indicate that most of the identified TFs are required for seedling root growth, with a subset affecting responses to CWD or hyperosmotic stress. Importantly, one of the three transcriptional regulators (*PIL2*, *MYB3R4*, *JMJ17*) that specifically affect stress responses in the growth assays, *JMJ17,* appears to be most relevant for regulatory processes, as it also affects CWD-induced JA and lignin production.

## Conclusion

To summarize, our aim was to identify transcriptional regulators that regulate primary cell wall metabolism as well as the response to CWD and hyper-osmotic stress in a THE1-dependent (and independent) manner. We have identified such regulators and established their involvement in stress responses and primary cell wall metabolism, providing starting points for analyzing the transcriptional regulation of primary cell wall metabolism and the response to CWD. The current estimate is that ∼10–15% of the ∼27k protein-coding genes in Arabidopsis (≈2,500–4,000) are involved in cell wall formation and maintenance (18, 19). While characteristic cell wall-related enzymes are easy to identify, relevant TFs, signal transduction elements, etc., are more difficult to isolate because cell walls are involved in many different biological processes. Previous work has shown that many TFs are required for secondary cell wall formation, illustrating the complexity of the transcriptional networks involved in cell wall metabolism (27). We think, therefore, the results presented here form a first small step towards dissecting the transcriptional network regulating primary cell wall formation.

### Plant material

The Arabidopsis Columbia ecotype (Col-0) was used as the wildtype. The Arabidopsis genotypes used in this study were obtained from the laboratories previously published or ordered from the Arabidopsis Biological Resource Center (ABRC) and are summarized in Supplemental Table S1. The obtained plants were propagated, and T-DNA insertions were verified by PCR using the Phire Plant Direct PCR Master Mix Kit (Thermo Fisher Scientific), followed by gel electrophoresis (1% agarose gel with SYBR Safe DNA Gel Stain (Thermo Fisher Scientific)). For primers, see Supplemental Table S2.

### Growth conditions

Arabidopsis seeds were grown under axenic conditions on half-strength Murashige Skoog medium supplemented with 1% sucrose (w/v, Sigma-Aldrich) and 0.05% MES (w/v, Sigma-Aldrich) under long-day conditions (16 h light at 22 °C / 8 h dark at 18 °C). For liquid cultures, seedlings were grown in 125 mL of growth medium in 250 mL Erlenmeyer flasks (Thermo Fisher Scientific) on an IKA KS501 flask shaker at 130 rpm. For hypersensitivity- and lignin deposition assays, seedlings were grown on solid medium (including 0.8% Gellan Gum (w/v, Thermo Fisher Scientific)).

### RNA isolation and sequencing

Col-0, *the1-1*, and *the1-4* seedlings were grown in liquid medium. Seven days after germination (7 DAG), seedlings were treated by exchanging the medium with one of the following conditions: 0.1% DMSO (v/v, mock), 600 nM ISX, 300 mM sorbitol, or a combination of 600 nM ISX and 300 mM sorbitol. After 2 hours, plant material was flash-frozen in liquid nitrogen and stored at −80 °C until further use. Total RNA was extracted using the Spectrum™ Plant Total RNA Kit combined with an on-column DNase I system (On-Column DNase I Digestion Set, both Sigma-Aldrich) to eliminate residual genomic DNA. RNA quality was assessed using the Agilent RNA 6000 Nano Kit with a Bioanalyzer (Agilent Technologies). RNA sequencing libraries were generated for 36 RNA samples using the SENSE mRNA-Seq library prep kit V2 (Lexogen GmbH). In brief, 600 ng of total RNA was incubated with magnetic beads coated with oligo-(dT) oligomers, and all RNAs except bound poly(A) RNA (mRNA) were removed by washing. Library preparation was then initiated by random hybridization of starter/stopper heterodimers containing Illumina-compatible linker sequences to the mRNA on the magnetic beads. A single-tube reverse transcription and ligation reaction extends the starter to the next hybridized heterodimer, where the newly synthesized cDNA insert was ligated to the stopper. Second-strand synthesis was performed to release the library from the beads. The resulting double-stranded library was purified and amplified (95°C, X min, 13 PCR cycles of [95°C x s, 68°C x s, 72°C x s], 72°C x min) after adding the adaptors and indexes. Finally, libraries were quantified by qPCR using Collibri™ Library Quantification Kit (Thermo Fisher Scientific) and validated using DNA High Sensitivity Reagent Kit on a Labchip GX (Revvity). The size range of the DNA fragments was measured to be in the range of 200-400 bp with an average library size of 244 bp. Libraries were normalized to 2.6 pM and clustered. Single-read sequencing was performed for 86 cycles (insert read) and 6 cycles (index read) on two NextSeq500 HO flow cells (Illumina), following the manufacturer’s instructions. FASTQ files were generated using bcl2fastq2 Conversion Software v2.20.0.422 (Illumina). Reads were quality-checked with FastQC and summarised with MultiQC. Adapter trimming and quality filtering were done with fastp v0.20.0. FastQ Screen confirmed that >97 % of reads matched the A. thaliana reference. Trimmed reads were aligned to the TAIR10 genome with STAR v2.7.8a using the AtRTD2 annotation. The mean mapping rate was 98 %, with 72–83 % of reads uniquely aligned. Gene-level counts were taken from STAR output (stranded forward). Quality metrics from Picard RnaSeqMetrics v2.23.4 indicated about 53 % mRNA bases and minimal 5′–3′ bias. Count data were analysed in R 3.6.1 using edgeR 3.26.8 and limma 3.40.6. Genes with CPM < 1 in fewer than two samples were excluded. Libraries were normalised by TMM and modelled with voom. Linear models were fitted per gene, and treatment effects were tested against mock controls. Differential expression was defined at FDR < 0.05 and |log_2_FC| > 0.585. The FASTQ files are available in the BioProject database (http://www.ncbi.nlm.nih.gov/bioproject) under accession PRJNA1260971.

### Quantitative analysis of transcript levels by qRT-PCR

Plant material (7 DAG) was harvested after a 6-hour treatment. Treatments were applied by exchanging the growth medium with one of the following conditions: 0.1% DMSO (v/v, mock), 600 nM ISX, 300 mM sorbitol, or a combination of 600 nM ISX and 300 mM sorbitol. Total RNA was extracted using the Spectrum™ Plant Total RNA Kit (Sigma-Aldrich) combined with an on-column DNase I system (On-Column DNase I Digestion Set; Sigma-Aldrich) to eliminate residual genomic DNA. RNA was quantified on NanoDrop One (Thermo Fisher Scientific), and 1,250 ng of total RNA was used for cDNA synthesis using the ImProm-II Reverse Transcription System (Promega). qRT-PCR was performed using a LightCycler 96 system (Roche) with SYBR Green I Master (Roche). Every sample was run in three independent biological and technical replicates. The expression data were normalized to the expression of *ACTIN2* (*ACT2*) and *EXOCYST COMPLEX COMPONENT SEC3A* (*SEC3a*). The primers used in RT-qPCR are listed in Supplemental Table S3.

### Hypersensitivity screen

Plants were grown on solid medium for 9 days under three conditions: control, 1.5 nM ISX, and 200 mM sorbitol. Plants were scanned and manually analysed in Fiji/ImageJ. Data visualization and statistics were performed in Prism 10 (GraphPad Software). Each genotype was analysed across three independent experiments, with 40 seedlings measured per experiment.

### Phytohormone analysis by UPLC-MS/MS

To assess phytohormone responses under stress conditions, seedlings were treated with sorbitol and isoxaben. Plants were grown in liquid media and treated at 7 DAG by exchanging the medium with one of the following conditions: 0.1% (v/v) DMSO (control), 600 nM ISX, 300 mM sorbitol, or a combination of 600 nM ISX and 300 mM sorbitol. Samples were collected at different time points: those treated with sorbitol and the combined ISX–sorbitol treatment were harvested after 4 hours, while samples treated with ISX alone were collected after 6 hours.

Jasmonic acid (JA) and abscisic acid (ABA) were extracted and analyzed using ultra-performance liquid chromatography with tandem mass spectrometry (Waters Acquity I Class UPLC system coupled to Xevo TQ-XS triple quadrupole mass spectrometer, both Waters Corporation, Milford, Massachusetts, U.S.) as described in reference (53), with minor modifications.

Briefly, freeze-dried seedlings were homogenized and extracted using a buffer containing 10% methanol, 1% acetic acid, 50 ng/mL jasmonic acid-d₅, and 10 ng/mL abscisic acid-d₆ (both from CDN Isotopes, Pointe-Claire, Canada). Chromatographic separation was performed on a Waters Cortecs C18 column (2.7 μm, 2.1 × 100 mm) using water (mobile phase A) and acetonitrile (mobile phase B), both containing 0.1% formic acid, at a flow rate of 0.4 mL/min. The linear gradient was programmed as follows: 0-1.2 min, 20% B; 1.2-4 min, ramp to 95% B; 4-5 min, hold at 95% B; 5-6 min, return to 20% B; 6-7 min, re-equilibration at 20% B. Quantification of jasmonic acid (JA) and abscisic acid (ABA) was performed using isotopic dilution calibration curves ranging from 0.2 to 1000 ng/mL. The limit of quantification (LOQ) for both analytes was 0.2 ng/mL. Each genotype was analyzed in at least three independent experiments in three technical replicates. The data were visualized and statistically validated using Prism 10 software.

### Sample preparation for cell wall characterization and cellulose quantification

Alcohol-insoluble residues (AIR) were prepared to enable cell wall characterization/analysis and cellulose quantification. Seedlings were harvested (7 DAG), flash-frozen in liquid nitrogen, and ground using a mortar and pestle. The powdered tissue was incubated at 70 °C for 30 min in 50 mM Tris buffer (pH 7.2) containing 1% SDS (w/v, Thermo Fisher Scientific) and centrifuged. The pellet was washed three times with water, with centrifugation following each wash. Samples were then incubated overnight at 65 °C in 80% ethanol. After pelleting, the ethanol was replaced with acetone, and samples were incubated for an additional 24 hours at 65 °C. The material was air-dried overnight and then incubated in 90% DMSO (Thermo Fisher Scientific) for 24 h. Next, another volume of 90% DMSO was added and the samples were incubated for another 24 h. Finally, the samples were washed three times with water, three times with 70% ethanol, and three times with acetone, and then air-dried.

### Updegraff crystalline cellulose quantification assay

The protocol was adapted from Updegraff (111) with minor modifications. The AIR samples (10-50 mg) were transferred into pre-weighed 2 mL microcentrifuge tubes. These were mixed with 1 mL of Updegraff reagent (acetic acid: nitric acid: water, 8:1:2 v/v; Thermo Fisher Scientific) and heated in an oven at 100 °C for 90 min. The sample was cooled to room temperature and then centrifuged at 10,000 x g for 5 min at room temperature to pellet the crystalline cellulose. The supernatant was removed, and the pellet was washed twice with 80% ethanol and twice with acetone at room temperature. After the acetone wash, the pellet was air-dried overnight at 40 °C. The samples were weighed, and 1 mL of 67% sulfuric acid (Thermo Fisher Scientific) was added to each 2 mL microcentrifuge tube. The tubes were then shaken on a rotary shaker at room temperature until all samples were dissolved. 20 μL of each sample was transferred to a glass tube and mixed with 1 mL of 0.3% anthrone (w/v, Thermo Fisher Scientific. Next, the samples were incubated at 100 °C for 5 min. The samples were then transferred to cuvettes, and the optical density (OD) was measured at 620 nm. Glucose concentrations were obtained by comparing sample OD to a standard regression equation.

### Cell wall monosaccharide composition analysis

For analysis of neutral cell wall sugars and uronic acids from non-crystalline cell wall matrix polymers, 1–2 mg AIR was weighed in 2-mL screw cap tubes and hydrolysed in 4% sulfuric acid (w/v) by autoclaving at 121°C for 60 min as described previously (112). Monosaccharide quantification was performed via high-performance anion-exchange chromatography with pulsed amperometric detection (HPAEC-PAD) using a biocompatible Knauer Azura HPLC system and an Antec Decade Elite SenCell detector as described previously (53). Monosaccharides were separated on a Thermo Fisher Dionex CarboPac PA20 column with a solvent gradient of (A) water, (B) 10 mM NaOH and (C) 700 mM NaOH at 0.4 mL/min flow rate: 0 to 25 min: 20% B (solvent % in A); 25 to 28 min: 20% to 0% B, 0% to 70% C; 28 to 33 min: 70% C; 33 to 35 min: 70% to 100% C; 35 to 38 min: 100% C; 38 to 42 min: 0% to 20% B, 100% to 0% C; 42 to 60 min: 20% B.

### Quantification of lignin deposition in Arabidopsis roots stained with phloroglucinol-HCl

Plants were grown on solid media for 7 DAG and placed in 6-well plates with mock conditions or 600 mM ISX. After 12 h, seedlings were collected in 70% ethanol, stained with 0.3% phloroglucinol (w/v: one volume of concentrated HCl (12M) to two volumes of 3% phloroglucinol in ethanol; Thermo Fisher Scientific), and imaged with a Zeiss Axio Zoom.V16 microscope, equipped with a PlanNeoFluar Z 1×/0.25 FWD 56 mm objective lens, a 25×/10 focal length eyepiece, and a Zeiss Axiocam506 color camera. Quantification of lignin deposition in the stained area of the root was measured using a Python script, thresholding the red coloration and calculating the fraction of coloured area in the root (n ≥ 15 roots per genotype). A one-way ANOVA was performed to evaluate the statistical significance of the data, followed by the Dunnett test.

## Acknowledgments

The work was supported by the European Commission under its Horizon 2021 framework program, Marie Sklodowska-Curie grant agreement (HORIZON-MSCA-2021-PF-01-01, Grant No. 101066093), the Research Council of Norway (NFR, Grant No. 315325), and the European Research Council (ERC, Grant No. 101118769). The library preparation and sequencing analysis were performed in close collaboration with the Genomics Core Facility (GCF) at the Norwegian University of Science and Technology (NTNU). GCF is funded by the Faculty of Medicine and Health Sciences at NTNU and the Central Norway Regional Health Authority. The authors gratefully acknowledge the support and access to mass spectrometry facilities provided by the NTNU Natural Science Faculty Mass Spectrometry Laboratory for their analysis of phytohormones. We thank Nadja Braun (Philipps-Universität Marburg, Germany) for technical assistance with cell wall monosaccharide analysis.

## Author Contributions

TT and TH designed the research. TT, SZ, TE, and ZB performed experiments and collected data. TT, SZ, TE, ZB, and LB analyzed the data. TT and TH prepared the figures and wrote the first draft. All authors discussed the results and edited the manuscript. TT and TH supervised the project and secured funding.

## Classification

Major: Plant biology; Minor: Stress biology

