## Supplemental Materials for "Coping with stress: Transcriptional regulators linking cell wall integrity maintenance with primary-wall metabolism in *Arabidopsis thaliana*"

A

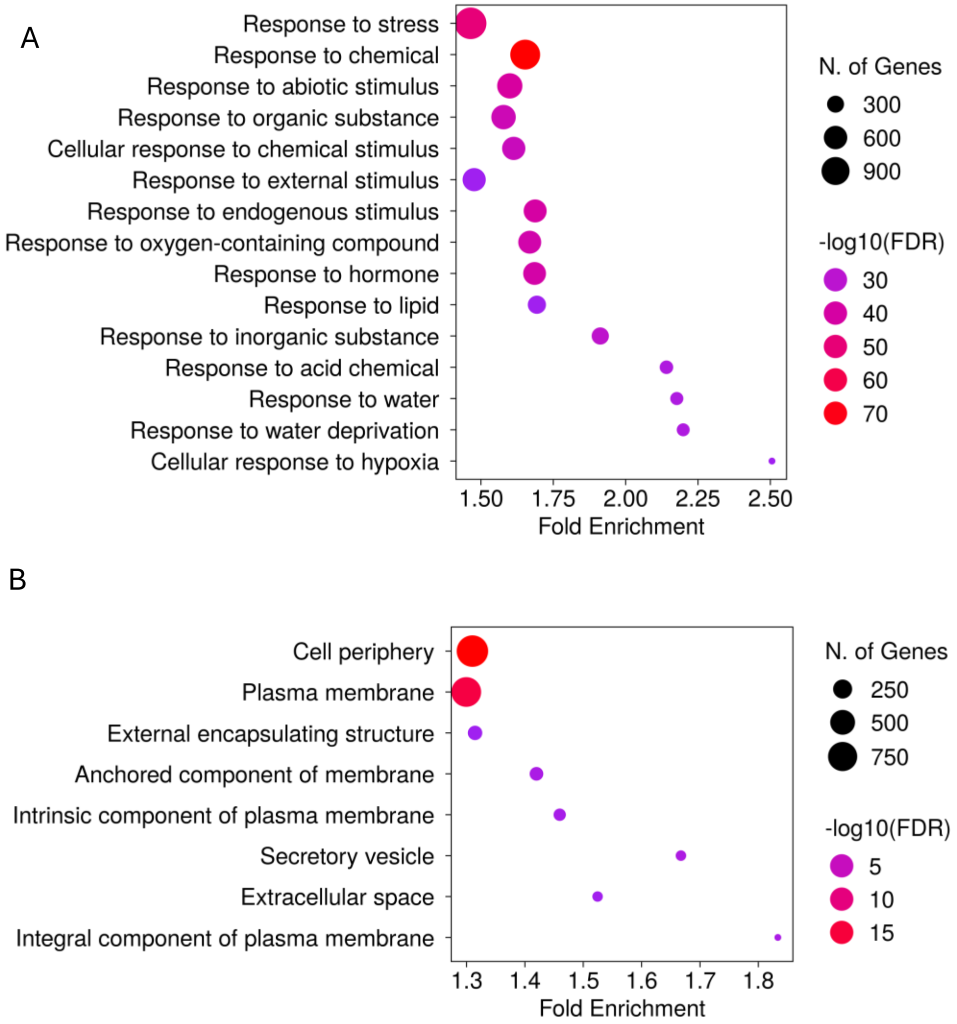


B



**Fig. S1. Gene Ontology (GO) enrichment analysis of differentially expressed genes (DEGs) in responseto stress conditions.**

GO enrichment analyses were performed for DEGs identified in Col-0, *the1-1*, and *the1-4* under Isoxaben (ISX), sorbitol (Sorb), and combined ISX + Sorb treatments. (A) GO enrichment for Biological Process. The top 15 GO terms (FDR < 0.05) are shown for each genotype. (B) GO enrichment for Cellular Component. The top 15 GO terms (FDR < 0.05) are shown for each genotype. The X-axis represents fold enrichment of each GO term. The size of each circle corresponds to the number of genes associated with that term, and the color gradient from purple to red indicates $-\log_{10}$(FDR), with red representing higher statistical significance. Analyses were performed using ShinyGO v0.82 (Ge SX, Jung D & Yao R, Bioinformatics 36:2628–2629, 2020).

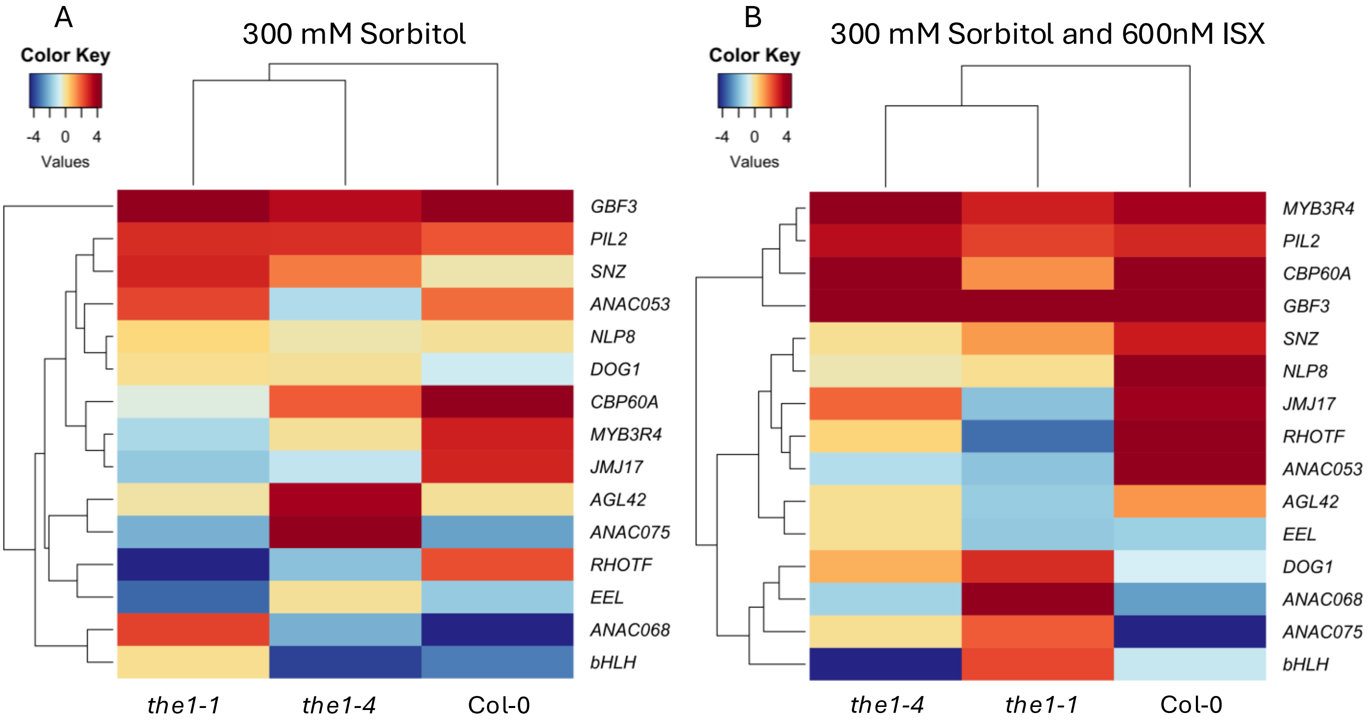


**Fig. S2. Heatmap analysis of candidate gene expression from RNAseq transcriptomic experiments in responce to osmotic and combined stress treatments.**

(A) Heatmap showing expression levels of selected candidate genes in Col-0, *the1-1*, and *the1-4* following treatment with 300 mM sorbitol. (B) Heatmap showing expression levels of the same candidate genes in Col-0, *the1-1*, and *the1-4* following combined treatment with 600 nM Isoxaben (ISX) and 300 mM sorbitol. Expression values are derived from RNA sequencing (RNAseq) transcriptomic experiments and normalized across genotypes and treatments. Color scaling reflects relative fold change with thresholds set between –4.5 and 4.5. Hierarchical clustering using Euclidean distance and complete linkage is shown as dendrograms for both rows and columns.

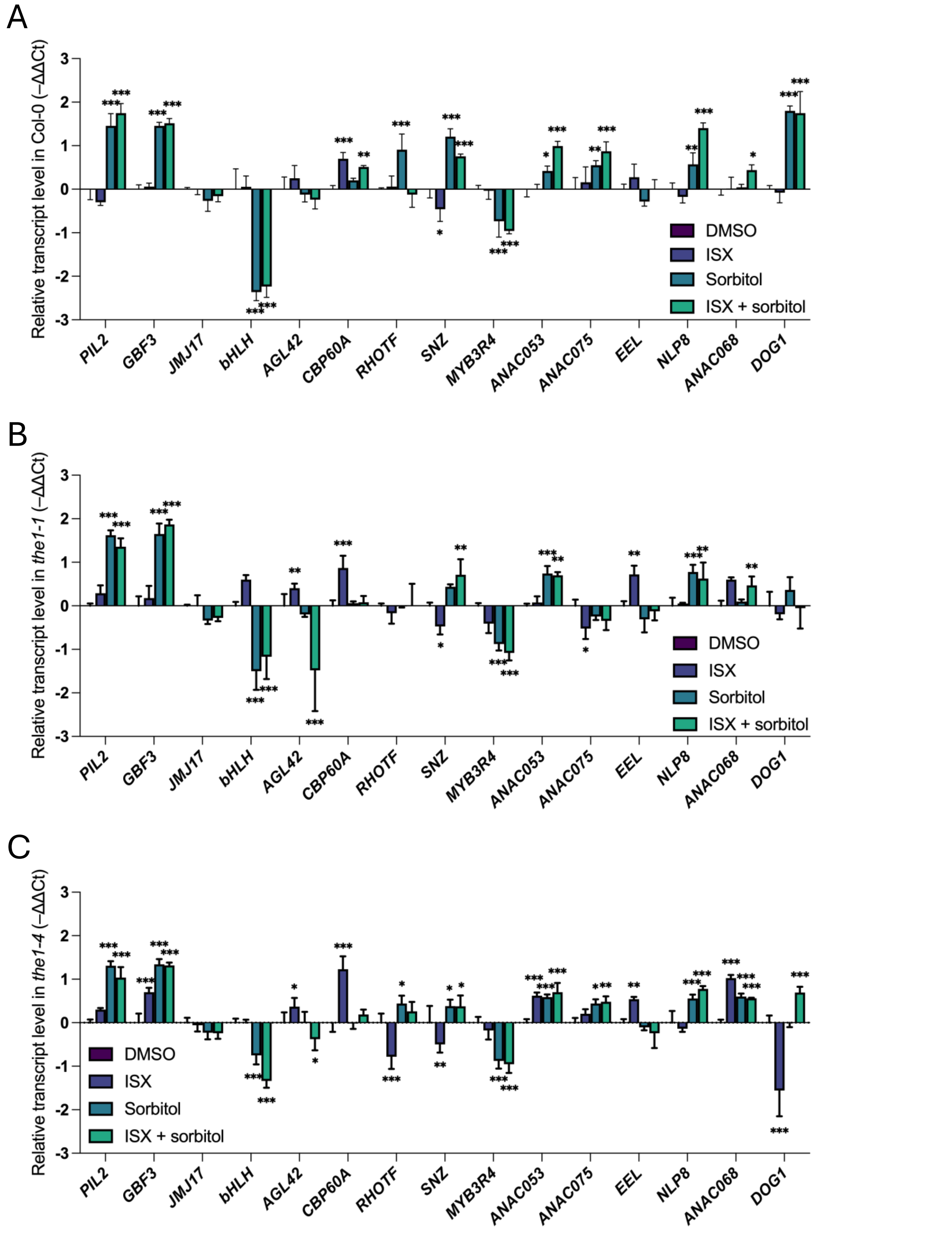


**Fig. S3. Quantitative RT-PCR analysis of candidate transcript levels in Col-0, *the1-1,* and *the1-4* seedlings in response to stress treatments.**

Expression levels of selected candidate genes in Arabidopsis seedlings after 6 h of treatment with control (DMSO), 600 nM Isoxaben (ISX), 300 mM sorbitol (Sorb), or combined ISX + Sorb. (A) Col-0 (B) *the1-1* (C) *the1-4*. Relative transcript level was analyzed by qRT-PCR and normalized to *SEC3a*. Data are presented as –ΔΔCt values ($\log_2$-transformed), calculated relative to DMSO-treated samples of each respective genotype. Bars represent mean ± SD (n ≥ 3). Asterisks indicate statistically significant differences compared to DMSO controls (two-way ANOVA followed by Dunnett's test; *$p < 0.05$; **$p < 0.01$; ***$p < 0.001$).

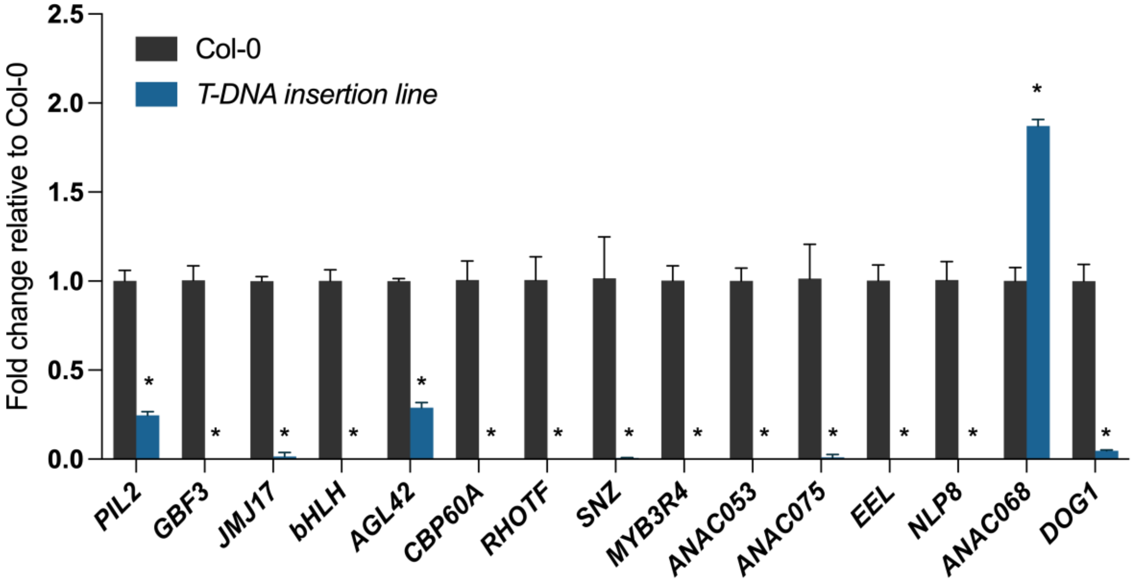


**Fig. S4. Quantification of candidate gene expression in T-DNA mutant lines.**
Expression levels of selected candidate genes in their corresponding T-DNA insertion mutant lines, shown as fold change relative to Col-0 mock. Transcript levels were normalized to *ACT2*. Data represent mean ± SD from 3 independent biological replicates. Asterisks indicate statistically significant differences compared to Col-0 (Student's t-test; $p < 0.05$).

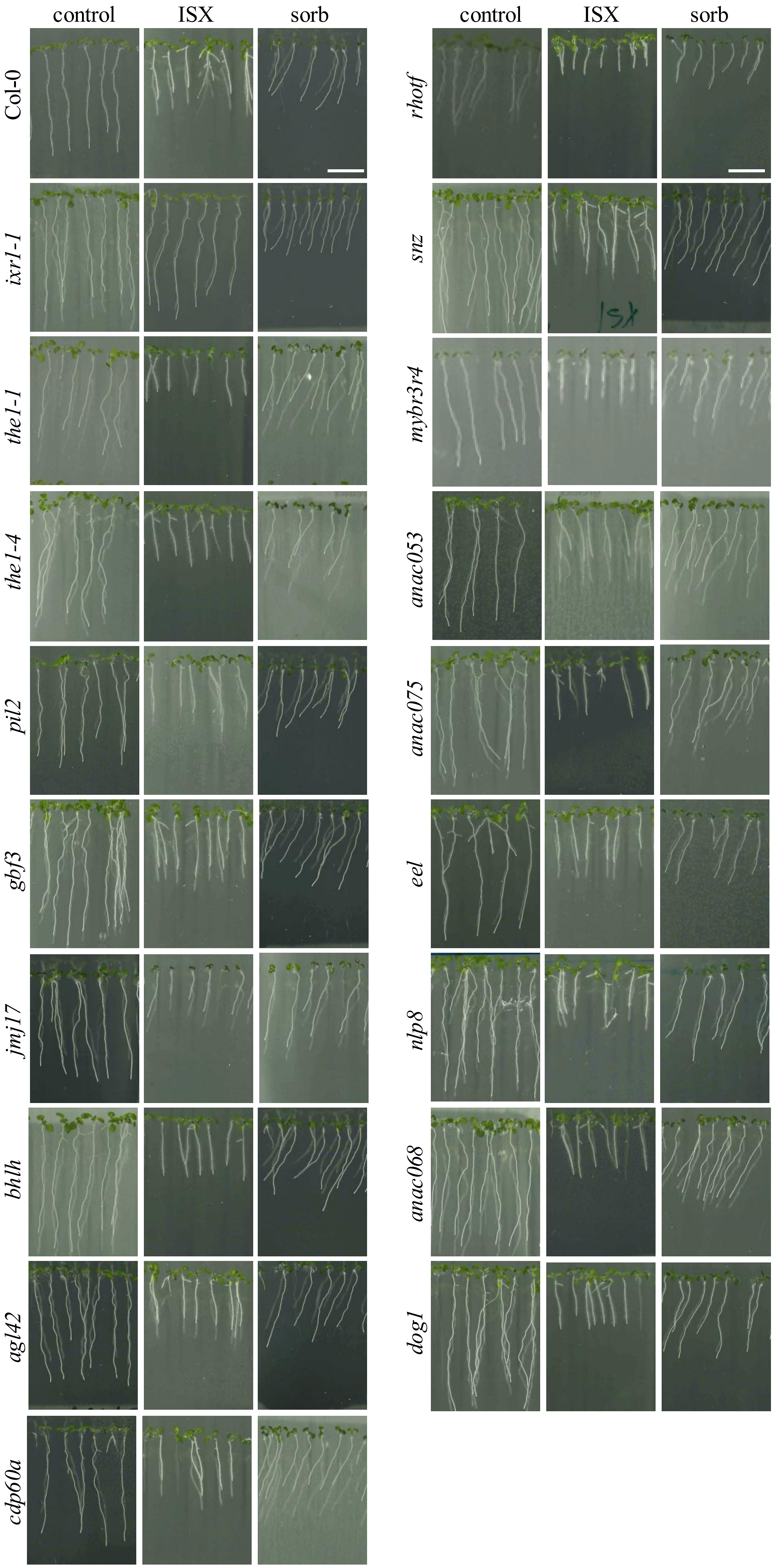


**Fig. S5. Representative images of Arabidopsis seedlings grown in control and stress conditions used in the hypersensitivity assay.** Representative images of Arabidopsis seedlings from the indicated genotypes grown in control conditions, 1.5 nM Isoxaben (ISX), and 200 mM sorbitol. Images were taken 7 days after germination (7 DAG). Scale bar: 1 cm.

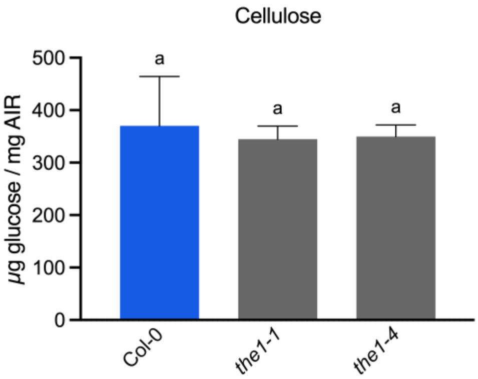


**Fig. S6. Cellulose content in *the1-1* and *the1-4* seedlings.**
Cellulose content (µg glucose / mg alcohol-insoluble residue (AIR)) was measured using the anthrone assay in 7-day-old seedlings of Col-0, *the1-1,* and *the1-4* genotypes. Data represent mean ± SD from five independent biological replicates. Statistical significance was determined using one-way ANOVA followed by Tukey's HSD test ($p < 0.05$). Different letters indicate statistically significant differences between genotypes.

| Monosaccharides | Negative regulation by | Positive regulation by |
|---|---|---|
| **Fucose** | *JMJ17, AGL42, EEL, NLP8* | - |
| **Galacturonic acid** | *ANAC075* | *JMJ17, EEL, NLP8, MYB3R4, RHOTF, DOG1* |
| **Glucose** | - | *EEL, PIL2, SNZ, MYB3R4, RHOTF, GBF3, bHLH, CBP60A, DOG1* |
| **Arabinose** | *EEL, MYB3R4* | - |
| **Rhamnose** | *PIL2, SNZ, MYB3R4, ANAC053, DOG1* | - |
| **Galactose** | *SNZ, MYB3R4, RHOTF, DOG1* | - |
| **Mannose** | *RHOTF, ANAC075* | - |
| **Xylose** | - | *RHOTF* |
| **Glucuronic acid** | - | - |

**Fig. S7. Summary of negative and positive regulation by indicated candidate genes on measured monosaccharides.**
"-" indicates no candidate genes identified for that regulatory category. Negative regulation indicates candidate genes whose mutation results in an increase in the monosaccharide content compared to Col-0, suggesting they normally act to repress its accumulation. Positive regulation indicates candidate genes whose mutation results in a decrease in the monosaccharide content compared to Col-0, suggesting they normally promote its accumulation. Monosaccharide levels were determined from alcohol-insoluble residue (AIR) in 7-day-old seedlings using standard sugar composition analysis.

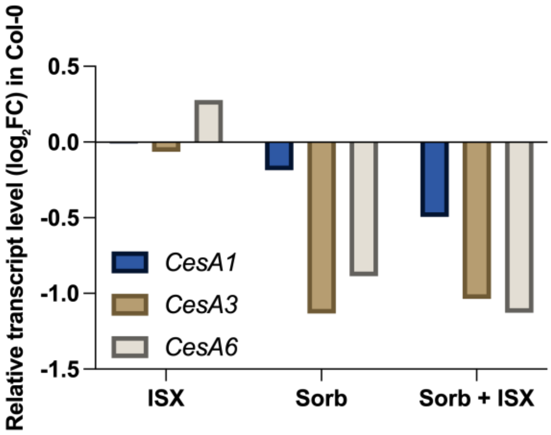


**Fig. S8. RNAseq-based transcript levels of primary cell wall cellulose synthases in response to ISX and osmotic stress.** The expression levels of *CesA1*, *CesA3*, and *CesA6* in Col-0 following in response to 2 hours of treatments with 600 nM ISX, 300 mM sorbitol, or their combination (600 nM ISX and 300mM sorbitol). Transcript abundance was quantified from RNA-seq–based transcriptomic data. Values represent $\log_2$ fold changes ($\log_2$FC) calculated from three biological replicates per treatment.

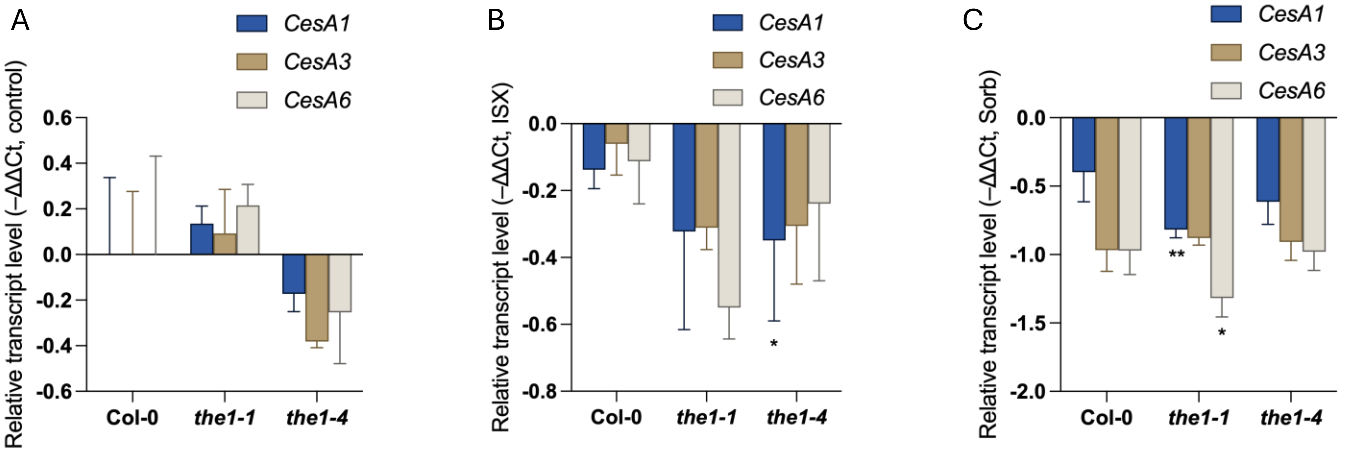


**Fig. S9. Relative transcript levels of *CesA1*, *CesA3*, and *CesA6* in Col-0, *the1-1*, and *the1-4* seedlings.**
Quantitative RT-PCR based analysis of *CesA1*, *CesA3*, and *CesA6* transcript levels in the indicated genotypes under (A) control conditions, (B) 600 nM Isoxaben (ISX), and (C) 300 mM sorbitol (Sorb) treatment. Transcript levels were normalized to *SEC3a* and are presented as –ΔΔCt values ($\log_2$-transformed relative expression). Bars represent mean ± SD of three independent biological replicates. Asterisks indicate statistically significant differences relative to Col-0 (two-way ANOVA followed by Dunnett's post hoc test; *p < 0.05; **p < 0.01; ***p < 0.001).

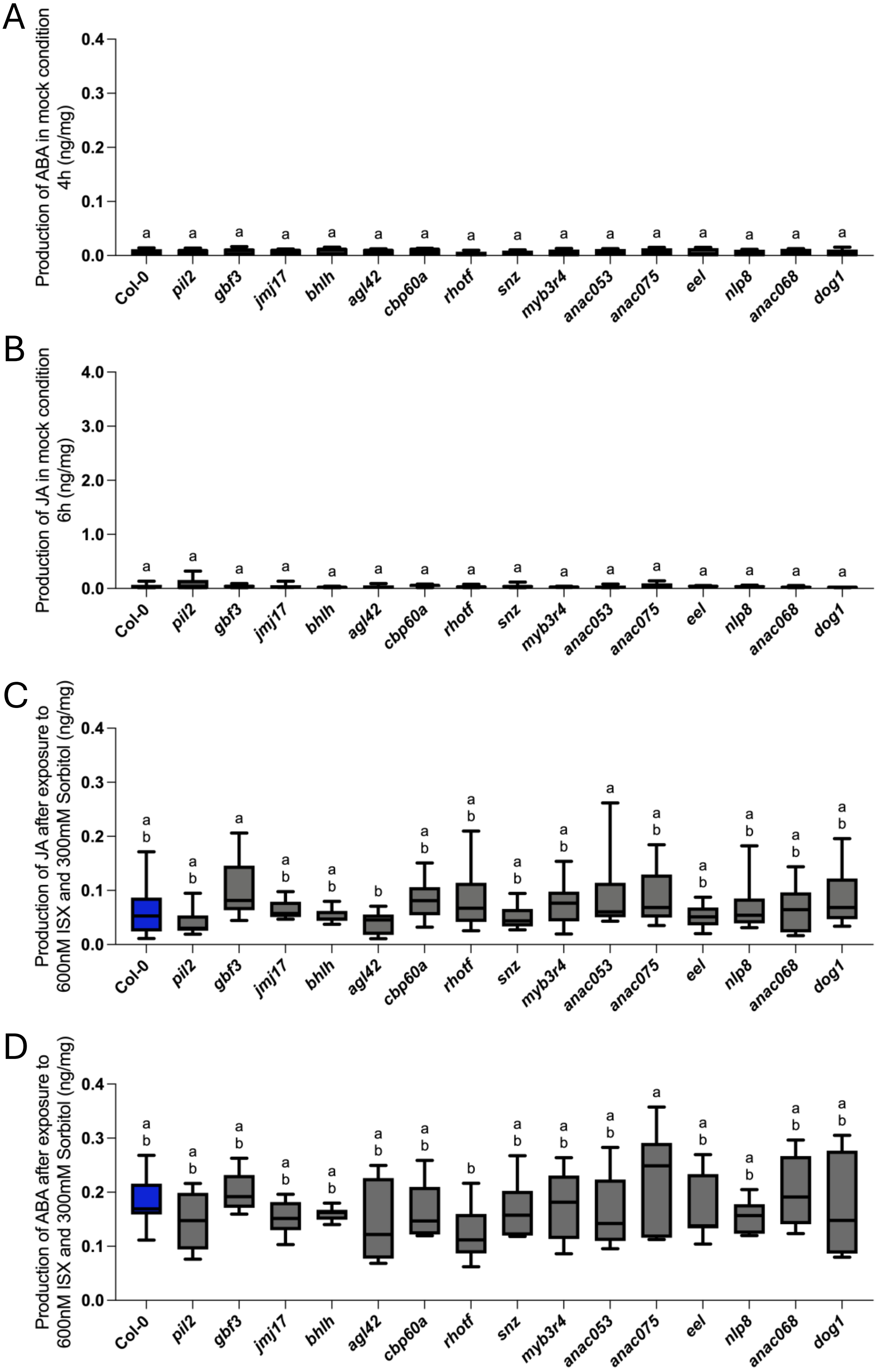


**Fig. S10. Jasmonic acid, abscisic acid phytohormone levels in candidate T-DNA insertion mutants under mock and combined stress treatments.**

(A) Abscisic acid levels (nanograms/milligrams dry weight) after 4 h in mock conditions (B) Jasmonic acid levels (nanograms/milligrams dry weight) after 6 h in mock (DMSO) conditions.. (C) Jasmonic acid levels (nanograms/milligrams dry weight) after 4 h of combined treatment with 600 nM Isoxaben (ISX) and 300 mM sorbitol. (D) Abscisic acid levels (nanograms/milligrams dry weight) after 4 h of combined ISX + sorbitol treatment. Graphs represent data from 3 to 4 independent experiments. Box plots display the full data range (min to max), with horizontal bars indicating the mean. Different letters denote statistically significant differences between genotypes (one-way ANOVA followed by Tukey's HSD test).

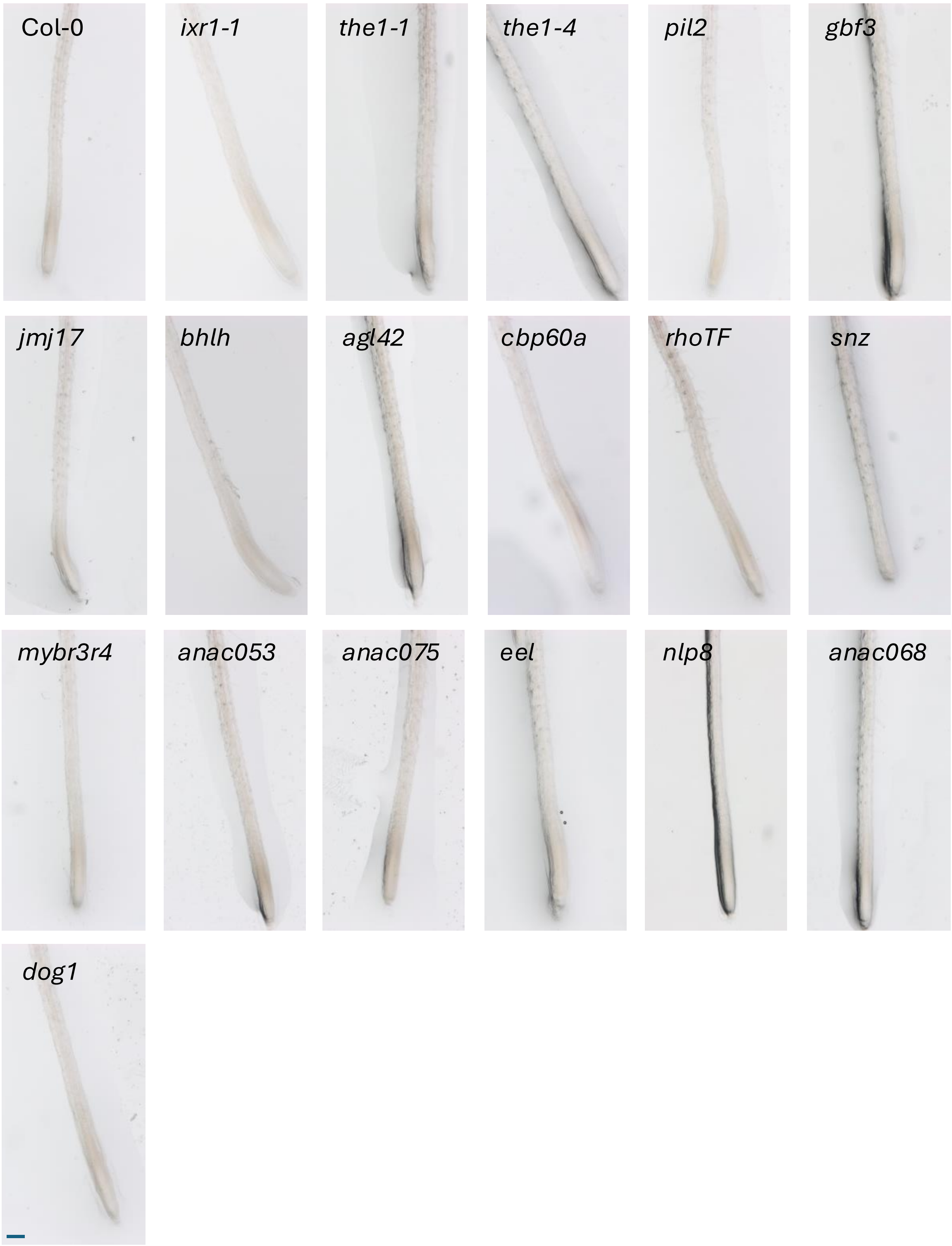


**Fig. S11. Representative images of lignin deposition in roots under control conditions.**
Representative images of seedlings from the indicated genotypes grown on ½ MS medium and transferred to mock treatment (fresh ½ MS medium) for 12 h. Roots were stained with phloroglucinol to visualize lignin deposition. Scale bar: 0.2 mm.

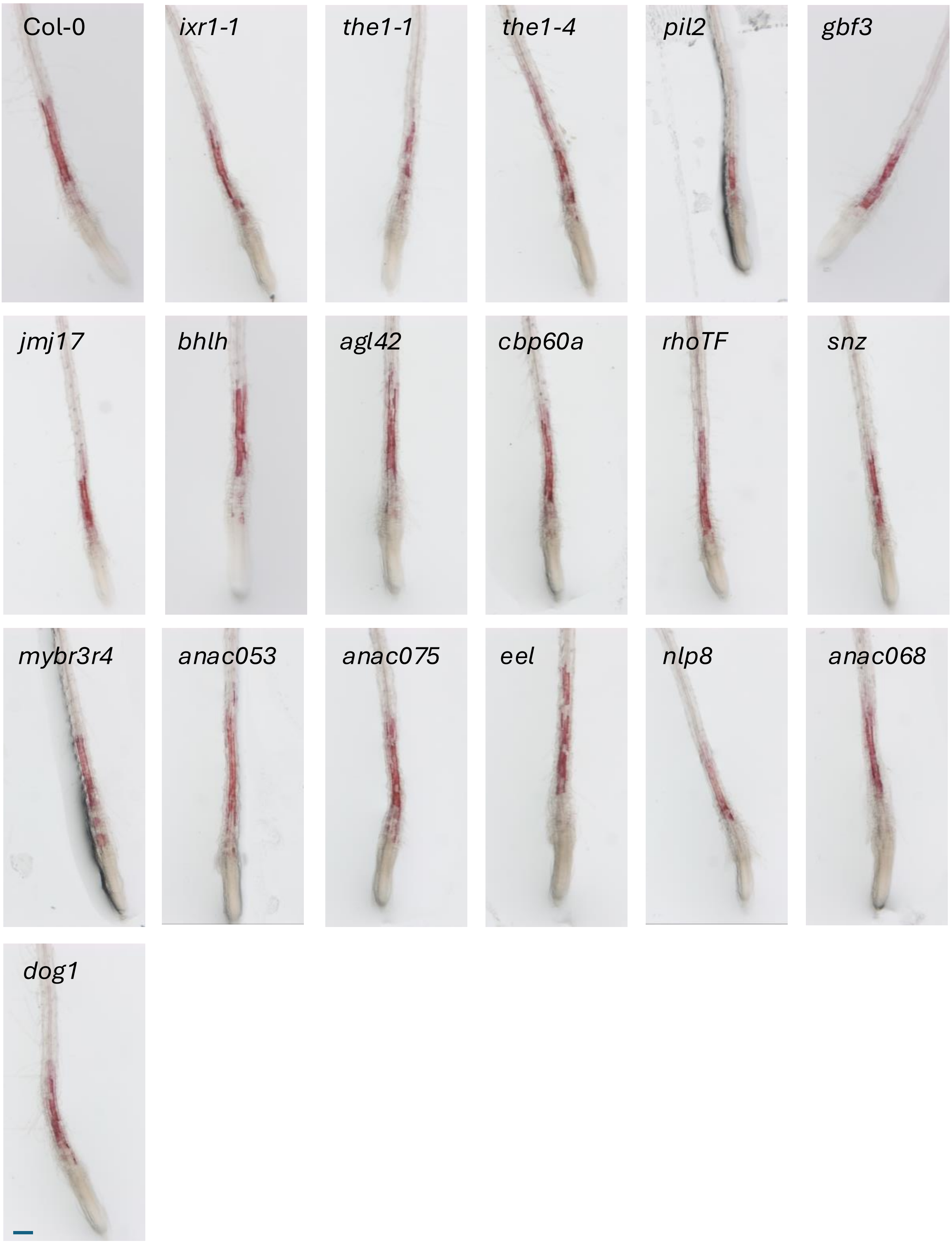


**Fig. S12. Representative images of lignin deposition in roots under ISX treatment.**
Representative images of seedlings from the indicated genotypes grown on ½ MS medium and treated with 600 nM Isoxaben (ISX) for 12 h. Lignin deposition in root tips was visualized using phloroglucinol staining. Scale bar: 0.2 mm.

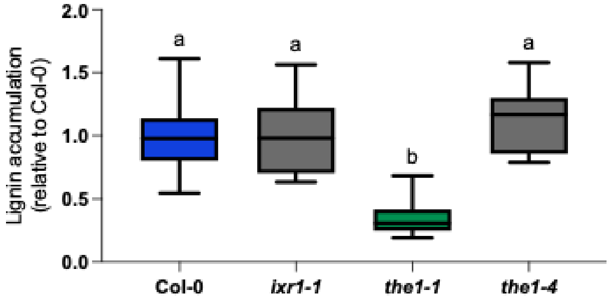


**Fig. S13. Lignin accumulation after 12 h of ISX treatment in the seedling.**
Relative lignin integrated density in root tips following 12 h of 600 nM Isoxaben (ISX) treatment, visualized by phloroglucinol staining and expressed as the ratio of mutant to Col-0. Data represent three independent experiments. Box plots display the full data range (min to max), with horizontal bars indicating the mean. Different letters denote statistically significant differences between genotypes (one-way ANOVA followed by Tukey's HSD test; $^*p < 0.05$).

Table S1: List of genotypes used in this study.

| Genotype | AGI | Description | reference |
|---|---|---|---|
| *ixr1-1* | AT5G05170 | CESA3 (Cellulose Synthase A3) ixr1-1 | Bacette |
| *the1-1* | AT5G54380 | THESEUS1 (THE1) | Hematy et al., 2007 |
| *the1-4* | AT5G54380 | THESEUS1 (THE1) | Mertz et al., 2017 |
| *pil2* | AT3G62090 | PHYTOCHROME INTERACTING FACTOR 3-LIKE 2; PIL2 (SALK_017311) | this paper |
| *gbf3* | AT2G46270 | G-BOX BINDING FACTOR 3; GBF3 (SALK_082840C) | (Ramegowda et al., 2017) |
| *jmj17* | AT1G63490 | JUMONJI DOMAIN-CONTAINING PROTEIN 17; JMJ17 (SALK_048786C) | this paper |
| *bHLH* | AT5G51780 | bHLH (SALK_087851C) | this paper |
| *agl42* | AT5G62165 | AGAMOUS-like 42; AGL42 (SALK_047915C) | this paper |
| *cbp60a* | AT5G62570 | CALMODULIN-BINDING PROTEIN 60A, CBP60A (SALK_036108C) | (Truman et al., 2013) |
| *rhotf* | AT4G18740 | Rho termination factor (SALK_038425C) | this paper |
| *snz* | AT2G39250 | SCHNARCHZAPFEN, SNZ (SALK_030031C) | (Mathieu et al., 2009) |
| *mybr3r4* | AT5G11510 | MYB DOMAIN PROTEIN 3R-4, MYB3R-4 (SALK_059819C) | (Haga et al., 2007) |
| *anac053* | AT3G10500 | NAC DOMAIN CONTAINING PROTEIN 53; NAC053 (SALK_009578) | N. P. Gladman, R. S. Marshall, K.-H. Lee, R. D. Vierstra, The Proteasome Stress Regulon Is Controlled by a Pair of NAC Transcription Factors in Arabidopsis. Plant Cell **28**, 1279–1296 (2016) |
| *anac075* | AT4G29230 | NAC DOMAIN CONTAINING PROTEIN 75, NAC075 (SALK_132120) | (Fujiwara and Mitsuda, 2016) |
| *eel* | AT2G41070 | ENHANCED EM LEVEL; EEL (SALK_021965) | (Ufaz et al., 2011) |
| *nlp8* | AT2G43500 | NIN-LIKE PROTEIN 8; NLP8 (SALK_031064) | (Yan et al., 2016) |
| *anac068* | AT4G01540 | ARABIDOPSIS NAC DOMAIN CONTAINING PROTEIN 68; ANAC068 (SALK_019893C) | this paper |
| *dog1* | AT5G45830 | DELAY OF GERMINATION 1, DOG1 (SALK_000867) | (Bentsink et al., 2006) |
